# Cooperativity of c-MYC with Krüppel-Like Factor 6 Splice Variant 1 induces phenotypic plasticity and promotes prostate cancer progression and metastasis

**DOI:** 10.1101/2024.01.30.577982

**Authors:** Sudeh Izadmehr, Heriberto Fernandez-Hernandez, Danica Wiredja, Alexander Kirschenbaum, Christine Lee-Poturalski, Peyman Tavassoli, Shen Yao, Daniela Schlatzer, Divya Hoon, Analisa Difeo, Alice C. Levine, Juan-Miguel Mosquera, Matthew D. Galsky, Carlos Cordon-Cardo, Goutham Narla

## Abstract

Metastasis remains a major cause of morbidity and mortality in men with prostate cancer, and the functional impact of the genetic alterations, alone or in combination, driving metastatic disease remains incompletely understood. The proto-oncogene c-MYC, commonly deregulated in prostate cancer. Transgenic expression of c-MYC is sufficient to drive the progression to prostatic intraepithelial neoplasia and ultimately to moderately differentiated localized primary tumors, however, c-MYC-driven tumors are unable to progress through the metastatic cascade, suggesting that a “second-hit” is necessary in the milieu of aberrant c-MYC-driven signaling. Here, we identified cooperativity between c-MYC and KLF6-SV1, an oncogenic splice variant of the KLF6 gene. Transgenic mice that co-expressed KLF6-SV1 and c-MYC developed progressive and metastatic prostate cancer with a histological and molecular phenotype like human prostate cancer. Silencing c-MYC expression significantly reduced tumor burden in these mice supporting the necessity for c-MYC in tumor maintenance. Unbiased global proteomic analysis of tumors from these mice revealed significantly enriched vimentin, a dedifferentiation and pro-metastatic marker, induced by KLF6-SV1. c-MYC-positive tumors were also significantly enriched for KLF6-SV1 in human prostate cancer specimens. Our findings provide evidence that KLF6-SV1 is an enhancer of c-MYC-driven prostate cancer progression and metastasis, and a correlated genetic event in human prostate cancer with potential translational significance.

## Introduction

Prostate cancer, the most diagnosed cancer in men, is a genetically heterogeneous and clinically complex disease (Siegel RL et al., 2021). Genetic alterations necessary or sufficient for prostate tumors to transition from localized to disseminated disease are incompletely understood. This gap in understanding has posed a challenge clinically in predicting tumor aggressiveness. Genomic and transcriptomic studies of human prostate tumors have identified commonly deregulated genetic events (Dhanasekaran SM et al., 2001, Lapointe J et al., 2004, Taylor BS et al., 2010, Grasso CS et al., 2012), many of which have since been functionally and mechanistically characterized using genetically engineered mouse models (GEMM). These prostate GEMMs have characteristically modeled the early stages of disease, but few have modeled the complete spectrum of disease development and progression.

Extensive scientific evidence has demonstrated that the activation of the cellular proto-oncogene c-MYC is a causative event in the induction of cancer and is frequently deregulated in human prostate cancer (Dhanasekaran SM et al., 2001, Lapointe J et al., 2004, Taylor BS et al., 2010, Grasso CS et al., 2012). *c-MYC* mRNA and nuclear c-MYC protein are expressed in early stages of human prostate cancer signifying that deregulated c-MYC protein is an early alteration in prostate tumorigenesis (Gurel B et al., 2008). Increased expression of c-MYC is sufficient to initiate prostatic intraepithelial neoplasia (PIN) and localized low-grade prostate adenocarcinoma *in vivo*, however, is insufficient to induce further progression suggesting that multiple genetic events are necessary and can cooperate to drive progression to metastatic disease (Zhang X et al., 2000, Ellwood-Yen K et al., 2003, Ellis L et al., 2016, Kim J et al., 2009, Yang G et al., 2012, Nguyen HG et al., 2018). Prior studies modeled by double transgenic or knock-out GEMMs of c-MYC and a “second-hit” (e.g., PTEN (Kim J et al., 2009), TMPRSS2-ERG (King JC et al., 2009), AKT (Clegg NJ et al., 2011), Caveolin-1 (Yang G et al., 2012), NF-κB (Jin RJ et al., 2008), PIM1 (Wang J et al., 2010), and Hepsin (Nandana S et al., 2010), Nanog (Liu B et al., 2017) have predominantly shown moderate degrees of cooperativity inducing mPIN and indolent disease, and in a few models (e.g. PTEN or RAS gain (Thompson TC et al., 1989, Hubbard GK et al., 2016, Arriaga JM et al., 2020)) progression was prompted to locally advanced and metastatic disease. GEMMs, which model human prostate cancer, can aid in our understanding of the role of c-MYC in cellular transformation and tumor progression. We previously identified splice variant 1 (SV1) of the KLF6 tumor suppressor gene, KLF6-SV1, as a metastasis-promoting late event in tumorigenesis that induces invasion and dedifferentiation in prostate cancer (Narla G et al., 2005; Narla G et al., 2005; Narla G et al., 2008). Due to splicing, KLF6-SV1, a cytoplasmic protein, lacks the C-terminus C2H2 three zinc-finger DNA binding domains, characteristic of Krüppel-like factors, and a nuclear localization signal (Narla G et al., 2005). A novel C-terminal sequence of 21 amino acids replaces these regions. Increased KLF6-SV1 expression has been associated with prostate cancer tumor growth, metastasis, hormone refractory disease, and poor survival (Narla G et al., 2005; Narla G et al., 2005; Narla G et al., 2008).

The multistep nature of tumorigenesis has been established to be in part due to the consecutive activation of several dominantly acting oncogenes. Here, we have taken two genetic alterations, KLF6-SV1 and c-MYC, observed in human prostate tumors that have temporal significance and modeled them in mice and human cells. We report for the first time a KLF6-SV1 GEMM and a double-transgenic GEMM expressing both c-MYC and KLF6-SV1, herein MYC-SV1, that faithfully recapitulate the molecular features and clinical phases of human prostate cancer. Coupled with supportive clinical and comprehensive proteomic studies, we define the oncogenic splice variant, KLF6-SV1, as a major contributor to c-MYC-driven prostate tumorigenesis and metastatic disease through the upregulation of vimentin. We demonstrate that disrupting the cooperativity of c-MYC and KLF6-SV1 can effectively prevent tumor progression. These findings provide further molecular and mechanistic insights into the oncogenic drivers of prostate cancer progression.

## Results

### KLF6-SV1 expression induces premalignant transformation in the mouse prostate

Our central hypothesis is that aberrant KLF6-SV1 expression in c-MYC-driven prostate cancer is sufficient to drive localized tumors through the full metastatic cascade. To test this hypothesis and determine whether KLF6-SV1 contributes to disease initiation *in vivo*, we engineered transgenic mice expressing the human KLF6-SV1 gene under the control of the β-Actin promoter like designs used previously in prostate mouse models (Roh M et al., 2006; Kim J et al., 2009) (Figure 1A). Prostates from KLF6-SV1 mice were harvested and compared to wild-type (WT) littermate controls (Figure 1B, C). Immunohistochemistry (IHC) and Hematoxylin and Eosin (H&E) staining were used to histologically evaluate the cells of the mouse prostate (Figure 1D-F). KLF6-SV1 GEMM harbored focal low-grade mouse prostatic intraepithelial neoplasia (mPIN) by 10 months of age (n=8/11 mice, *P* < 0.0001), suggesting that KLF6-SV1 can induce pre-neoplastic transformation. A subset of KLF6-SV1 GEMM developed benign hyperplasia of the prostate (*n* = 2/11 mice) or remained unchanged (*n* = 1/11) (Figure 1E, F). mPIN lesions were in the dorsal (DP) and anterior (AP) prostate lobes of the mouse prostate. α-SMA and p63 IHC staining confirmed the *in situ* nature of the mPIN lesions in KLF6-SV1 GEMM by the presence of an intact basal layer (p63-positive basal cells) and fibromuscular layer (α-SMA-positive fibromuscular cells) (Figure S2B, S2C). However, the mPIN lesions were self-limited during the lifespan of the mice. KLF6-SV1 expression alone was insufficient to initiate an overt tumor of the mouse prostate in the specific genetic context of this model, providing evidence that KLF6-SV1 is sufficient to induce a pre-cancer phenotype in the prostate modeling the early stages of human prostate cancer similar to those observed in other established transgenic models of the disease (e.g. AKT, TMPRSS2-ERG, PTEN, and c-MYC (Zhang X et al., 2000; Ellwood-Yen K et al., 2003; King JC et al., 2009; Clegg NJ et al., 2011)).

**Figure 1.**
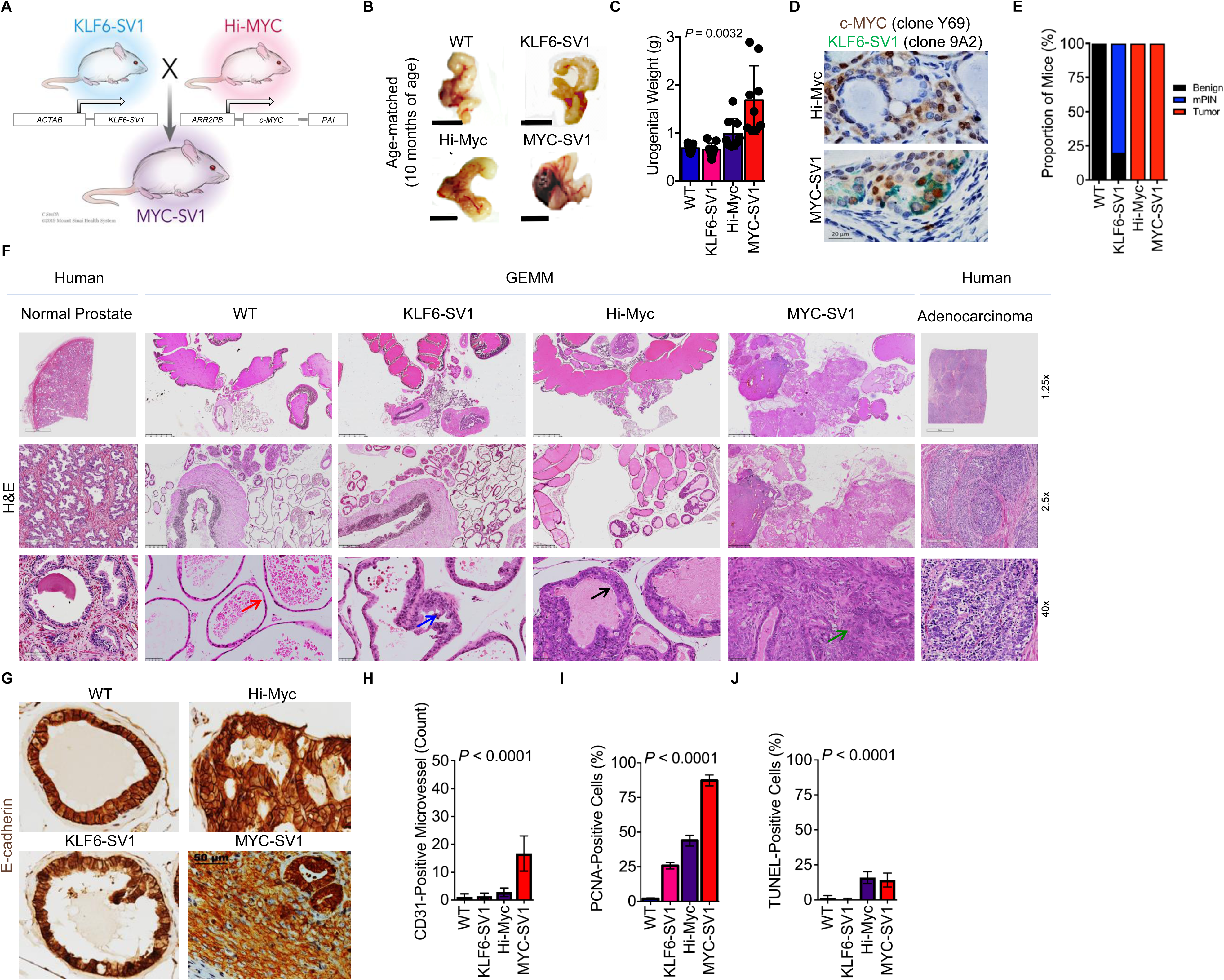
KLF6-SV1 cooperates with c-MYC to promote the progression of prostate tumorigenesis *in vivo*. **A**, Generation of MYC-SV1 GEMM. Confirmation of germline transmission of transgenes. **B,** Representative gross images of the urogenital system of WT, KLF6-SV1, Hi-Myc and MYC-SV1 mice at 10 months of age (scale bar=1cm). **C**, Urogenital weights of WT, KLF6-SV1, Hi-Myc and MYC-SV1 mice at 10 months of age (mean ± s.d., *P* = 0.0032, ANOVA). **D,** IHC staining of c-MYC (brown) and KLF6-SV1 (green) co-expression of FFPE sections of representative Hi-Myc and MYC-SV1 mouse prostates at 10 months of age (63x magnification; scale bar=20 µm). **E,** Penetrance of histological phenotype of WT (*n* = 10), KLF6-SV1 (*n* = 11), Hi-Myc (*n* = 10) and MYC-SV1 (*n* = 10) at 10 months of age. **F,** Comparative pathological characterization of the mouse prostate. Photomicrographs of FFPE hematoxylin and eosin (H&E) stained sections of representative WT, KLF6-SV1, Hi-Myc and MYC-SV1 prostates at 10 months of age of age, and paired normal and adenocarcinoma (grade IV) of human prostate tissue at low (1.25x magnification, scale bar=2.5mm; 2.5x magnification, scale bar=500µm) and high (40x magnification; scale bar=50µm) magnification views of the prostate are shown. WT mice display healthy glands (red arrow); KLF6-SV1 mice exhibit PIN (blue arrow); Hi-Myc mice develop focally invasive well-differentiated prostate cancer (black arrow); MYC-SV1 mice develop diffuse poorly differentiated invasive prostate cancer (green arrow). **G,** IHC staining of E-cadherin (brown) of FFPE sections from representative GEMM prostates at 10 months of age (40x magnification; scale bar=50 µm). **H,** Microvessel count and representative IHC of CD31 stained vessels per field (mean ± s.d., *P* < 0.0001, ANOVA); **I,** PCNA-positive proliferating cells per field (mean ± s.d., *P* < 0.0001, ANOVA); **J,** TUNEL-positive apoptotic cells per field (mean ± s.d., *P* < 0.0001, ANOVA)

### KLF6-SV1 drives the rapid progression of prostate adenocarcinoma in cooperation with c-MYC

To investigate whether KLF6-SV1 functions as a secondary genetic event to promote tumor progression in c-MYC-driven prostate cancer, we crossed KLF6-SV1 transgenic mice with Hi-Myc transgenic mice, a prostate-specific model that uses the epithelial-specific rat probasin promoter to overexpress human c-MYC (Ellwood-Yen K et al., 2003), to generate a novel double transgenic mouse model which we termed MYC-SV1 (Figure 1A). MYC-SV1 double transgenic mice displayed gross enlargement of the prostate (Figure 1B) and a significant increase in the weight of the urogenital system compared to WT, KLF6-SV1 or Hi-Myc mice (*P* < 0.0001, Figure 1C). Expression of human KLF6-SV1 and c-MYC protein in the GEMMs were confirmed by genotyping and IHC (Figure 1D). In line with previous reports, we found that Hi-Myc mice developed mPIN and then focal invasive well and moderately differentiated adenocarcinoma (Figure 1E, F) (Ellwood-Yen K et al., 2003). MYC-SV1 mice gave rise to overt, diffuse, and poorly differentiated invasive tumors in all lobes of the prostate (VP, DLP, and AP lobes). Histopathological analysis by H&E (Figure 1F, S1A, S1B, S1C, S4B, S4C), E-cadherin (Figure 1F), p63 (Figure S2B), and α-SMA (Figure S1C) immunohistochemistry staining showed that MYC-SV1 mice reliably developed highly penetrant mPIN, well and moderately differentiated adenocarcinoma, and then poorly differentiated invasive prostatic carcinoma. Tumor cells were positive for E-cadherin protein expression, confirming the epithelial origin of the lesions, with decreased expression of E-cadherin protein in poorly differentiated regions within the tumor (Figure 1G, S2C). MYC-SV1 prostate tumors were highly invasive and penetrated the basal layer, fibromuscular layer, stroma, periprostatic adipose tissue, and the capsule of the prostate (Figure S1B, S1C, S2B). Loss of the basal and fibromuscular layer of the prostate glands by the displacement of basal cells (p63-positive cells) and fibromuscular cells (α-SMA-positive cells), respectively, confirmed the invasive nature of the tumor cells (Figure S1C, S2B). Tumors displayed features of angiogenesis noted by increased blood vessel formation within the lesions (*P* < 0.0001, Figure 1H), were highly proliferative (*P* < 0.0001, Figure 1I) and apoptotic (*P* < 0.0001, Figure 1J).

### KLF6-SV1 and c-MYC promote tumor progression and metastasis *in vivo*

We followed a cohort of 93 mice over their lifetime for survival analysis. Consistent with the aggressive phenotype noted, a significant reduction in survival was observed in the MYC-SV1 mice compared with WT, KLF6-SV1, and Hi-Myc mice, which had similar lifespans (*P* < 0.0001, Figure 2A). We determined the kinetics of mPIN, tumor development, and progression of disease (Figure S4A-C). By 1 month of age, MYC-SV1 mice showed clear histological evidence of mPIN (*n* = 3/3) (Figure S4B, S4C). The latency of tumor development was shortened through the cooperativity of c-MYC and KLF6-SV1. Transition from mPIN to invasive carcinoma, and penetration of tumor cells into the surrounding stroma, was evident by 3 months of age. Mice developed pan-cytokeratin (Pan-CK) positive adenocarcinoma (*n* = 4/4) with invasive features (*n* = 2/4) (Figure S4B-D). In line with previous reports, only mPIN was found in Hi-Myc mice at this age (Ellwood-Yen K et al., 2003, Melis M et al., 2017). By 6 months of age, tumors progressed to high-grade poorly differentiated carcinoma (i.e. tumors of epithelial origin lacking glandular structure) (Figure S4C). Prostate lesions in Hi-Myc and KLF6-SV1 mice did not progress to advanced disease. Notably, aged MYC-SV1 mice developed large prostate tumors with cystic degeneration and bladder outlet obstruction causing hydronephrosis (Figure S5). Taken together these results suggest that KLF6-SV1 induces invasive and proliferative prostate carcinoma in the setting of c-MYC dysregulation.

**Figure 2.**
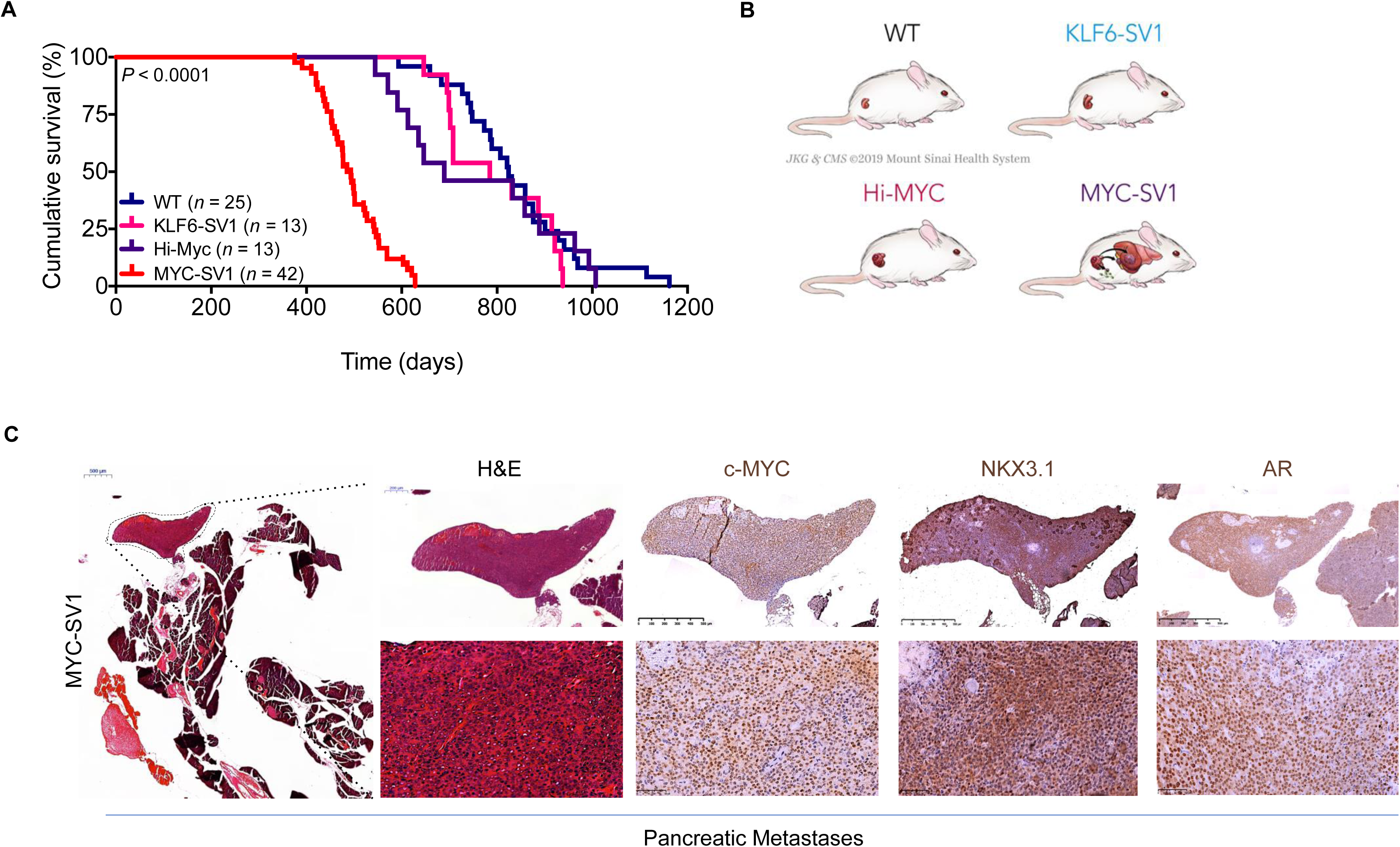
MYC-SV1 mice develop poorly differentiated tumors of luminal epithelial origin with metastatic disease. **A**, Kaplan–Meier cumulative survival analysis shows a significant decrease in the lifespan of MYC-SV1 mice (*n* = 42, Median survival 477 days) compared with WT (*n* = 25, Medial survival = 824 days), KLF6-SV1 (*n* = 13, Median survival = 785 days), and Hi-Myc (*n* = 13, Median survival = 689 days) cohorts (*P* < 0.0001, Log-rank test). **B**, Schematic of WT and GEMM metastatic phenotypes. **C,** Photomicrographs of H&E and immunohistochemical stained sections of MYC-SV1 metastatic pancreatic lesion at low and high magnification (2x magnification, scale bar=500µm; 40x magnification, scale bar=50µm) magnification. Metastatic tumor nodules encircled by dashed lines (2x magnification, scale bar=500µm). Pancreatic metastasis was stained for AR, c-MYC and NKX3.1 to confirm prostate origin.

Metastasis is a late-stage adverse health event in human prostate cancer (Siegel RL et al., 2021). Metastatic lesions were observed in MYC-SV1 mice at several anatomical sites including lymph nodes, liver, and pancreas (Figure 2B, 2C, S3, S6). A representative pancreatic lesion is shown in Figure 2C and S3 from a MYC-SV1 mouse at 12 months of age. Cell morphology on H&E, and positive AR, NKX3.1, and c-MYC protein expression confirmed the prostatic origin of the metastatic foci (Figure 2C). NKX3.1 protein expression is restricted to prostate epithelial cells and used as a diagnostic biomarker for prostate cancer and metastatic lesions originating from the prostate. Additionally, c-MYC is driven by the probasin promoter, which limits the expression of c-MYC to luminal epithelial cells. Metastatic foci also expressed diffuse cytoplasmic expression of E-cadherin and pan-cytokeratin (Figure S3). Lack of α-SMA or collagen protein expression (Masson’s Trichrome staining) confirmed that the metastatic foci were not of myoepithelial or fibroblast origin, respectively (Figure S3). Hi-Myc mice did not develop metastatic disease, a constrained phenotype that aligns with previous reports of the model (Ellwood-Yen K et al., 2003; King JC et al., 2009). MYC-SV1 mice modeled each phase of human prostate cancer development and progression, including mPIN, adenocarcinoma, and metastases. Overall, these data indicate that KLF6-SV1 is sufficient, in the presence of c-MYC, to drive the multi-step process of the metastatic cascade of prostate tumorigenesis.

### KLF6-SV1 and c-MYC expression is mutually regulated and significantly associated with proliferation

Our finding that cooperativity exists between KLF6-SV1 and c-MYC *in vivo* prompted us to investigate this relationship *in vitro* using a panel of human prostate cell lines spanning the spectrum of prostate cancer disease states. The expression of *KLF6-SV1* and *c-MYC* mRNA was measured in the human prostate cell lines (Figure 3A, B). There was a significant difference in mRNA (*c-MYC, P* < 0.001; *KLF6-SV1, P* = 0.0289) and protein levels amongst the non-tumorigenic and cancer cells. Increased c-MYC and KLF6-SV1 protein expression was observed in metastatic prostate cancer cells compared to PIN and non-tumorigenic prostate cells (Figure 3C). We next impaired the function of these genes by small interfering RNA (siRNA) individually or in combination in PC3 and PC3M cells, which harbor c-MYC amplification (Taylor BS et al. 2010) (Figure 3A, C), to evaluate the effects of c-MYC or KLF6-SV1 loss-of-function. Knockdown of KLF6-SV1 by siRNA in PC3 cells significantly reduced *c-MYC* mRNA and c-MYC protein expression within 48 hours (*P* < 0.001) (Figure 3D, E). Knockdown of c-MYC was associated with significantly reduced *KLF6-SV1* mRNA and KLF6-SV1 protein expression in PC3 cells. Similar changes in expression were observed in PC3M cells (Figure 3F, G). To complement the loss-of-function studies, we engineered stable cell lines which exogenously expressed c-MYC, KLF6-SV1, or both, to conduct gain-of-function studies in the benign luminal epithelial cell line, RWPE-1, that endogenously expresses low basal levels of c-MYC and KLF6-SV1 (Figure 3A-C). We again observed that endogenous expression of *KLF6-SV1* mRNA was significantly associated with an increase in c-MYC mRNA (*P* < 0.001), and KLF6-SV1 protein expression was associated with c-MYC protein expression (Figure 3H). Similarly, the added expression of c-MYC was associated with increased KLF6-SV1 mRNA and KLF6-SV1 protein expression (*P* < 0.001) (Figure, 3I-L). Next, we examined the effects of c-MYC and KLF6-SV1 on proliferation (Figure 3M-O) and the cell cycle (Figure 3P). Dual knockdown of c-MYC and KLF6-SV1 in PC3 cells also correlated with inhibition of cell proliferation demonstrated by a significant decrease in thymidine incorporation (*P* < 0.001) (Figure 3M) and KI-67 mRNA expression (*P* < 0.001) (Figure 3O). Additionally, silencing of c-MYC and KLF6-SV1 induced a cell cycle arrest in the G1/S phase with increased expression of CDKN1A (p21) (*P* < 0.0089) suggesting inhibition of cell growth and DNA replication (Figure 3Q, R).

**Figure 3.**
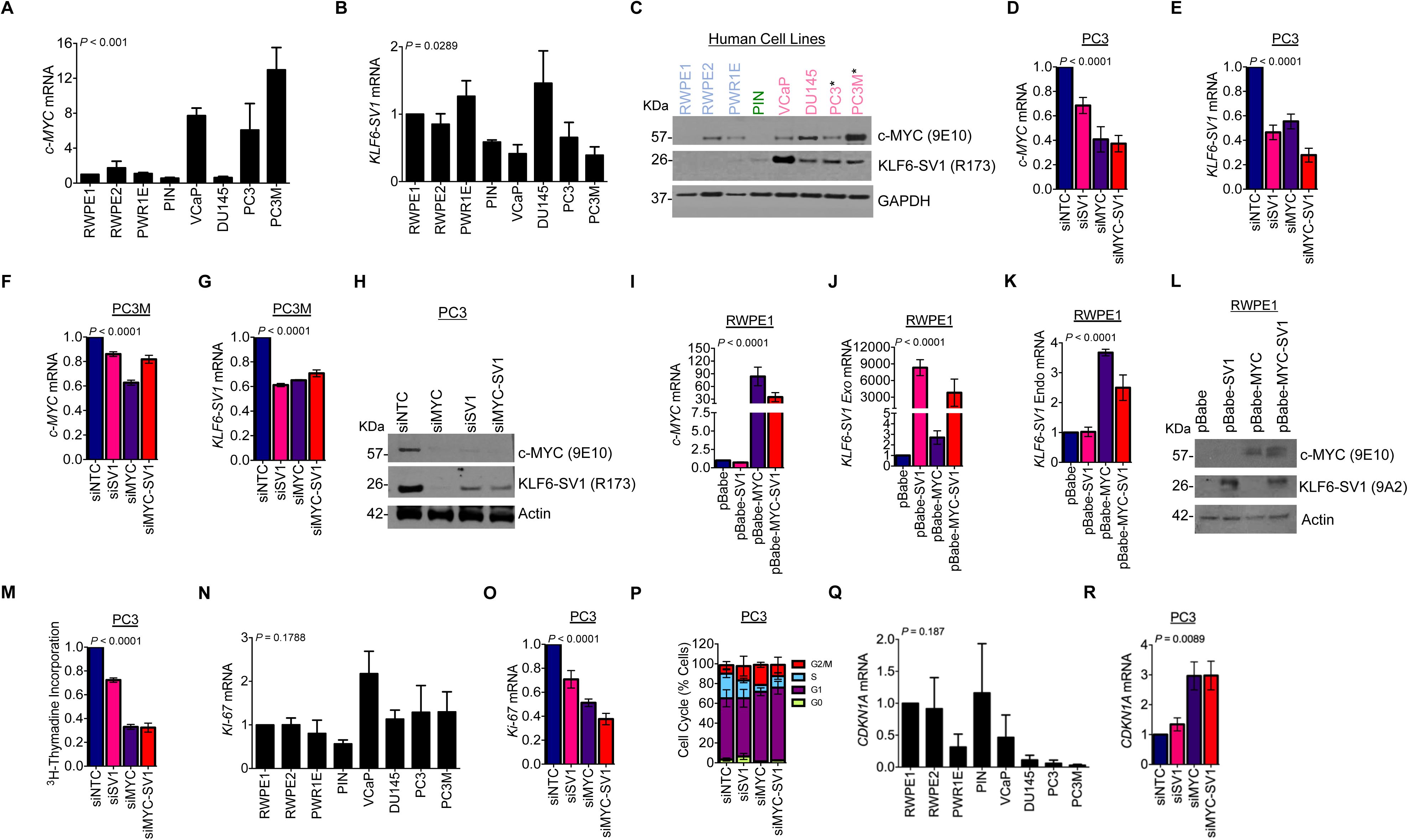
c-MYC and KLF6-SV1 cooperate and regulate cellular proliferation *in vitro*. **A**, *c-MYC* mRNA in a panel of prostate cell lines of differing disease states. Asterisk (*) denotes cell lines with c-MYC amplification (mean ± s.d., *P* < 0.001, ANOVA). **B,** *KLF6-SV1* mRNA in a panel of prostate cell lines of differing disease states (mean ± s.d., *P* = 0.0289, ANOVA). **C,** c-MYC and KLF6-SV1 protein expression in a panel of prostate cell lines of differing disease states (blue = benign cell lines, green = PIN cell lines, pink = tumorigenic cell lines; PC3 and PC3M cell lines with *c-MYC* amplification). **D,** qRT-PCR for c-MYC and **E**, *KLF6-SV1* mRNA expression 48 hours after *KLF6-SV1* and *c-MYC* siRNA-mediated knockdown in PC3 cells (mean ± s.d., *P* < 0.0001, ANOVA). **F,** qRT-PCR for c-MYC **G**, *KLF6-SV1* mRNA expression with KLF6-SV1 and c-MYC knockdown by siRNA in PC3M cells (mean ± s.d., *P* < 0.0001, ANOVA). PC3 and PC3M cells transfected with indicated siRNAs normalized to the corresponding non-silencing control. **H,** KLF6-SV1 and c-MYC protein expression 48 hours after KLF6-SV1 and c-MYC knockdown by siRNA in PC3 cells. **I**, qRT-PCR for c-MYC mRNA expression; **J**, qRT-PCR for KLF6-SV1 mRNA; **K**, qRT-PCR for exogenous KLF6-SV1 mRNA of RWPE-1 cell lines stably overexpressing p-Babe-KLF6-SV1, p-Babe-c-MYC, or p-Babe-KLF6-SV1/c-MYC in RWPE-1 cells (mean ± s.d., *P* < 0.0001, ANOVA). RWPE-1 cells transfected with indicated plasmids normalized to the corresponding p-Babe control. **L**, KLF6-SV1 and c-MYC protein expression of RWPE-1 cell lines stably expressing p-Babe-KLF6-SV1, c-MYC, or KLF6-SV1/c-MYC. **M**, *KI-67* mRNA expression in a panel of prostate cell lines of differing disease states (mean ± s.d., *P* = 0.1788, ANOVA). **N**, *KI-67* mRNA expression 48 hours after KLF6-SV1 and c-MYC knockdown by siRNA in PC3 cells (mean ± s.d., *P* < 0.0001, ANOVA). **O**, Thymidine incorporation 48 hours after KLF6-SV1 and c-MYC knockdown by siRNA in PC3 cells (mean ± s.d., *P* < 0.0001, ANOVA). **P**, Cell cycle analysis of PC3 cells 48 hours after KLF6-SV1 and c-MYC knockdown by siRNA. Bars in graphs represent three biological replicates. **Q,** *CDKN1A* (p21) mRNA expression in a panel of prostate cell lines of differing disease states (mean ± s.d., *P* = 0.187, ANOVA). **R,** *CDKN1A* (p21) mRNA expression 48 hours after KLF6-SV1 and c-MYC knockdown by siRNA in PC3 cells (mean ± s.d., *P* = 0.0089, ANOVA).

### Targeting of c-MYC through androgen ablation impairs tumor maintenance *in vivo*

Prior *in vivo* studies have demonstrated that c-MYC is necessary and sufficient for tumor maintenance (Podsypanina K et al., 2008; Soucek L et al., 2008). Since we observed that in PC3 cells with siRNA mediated silencing of c-MYC resulted in rapid growth arrest (Figure 3M, O), we uncoupled c-MYC and KLF6-SV1 expressions *in vivo* to study the effects of c-MYC depletion in the presence of functional KLF6-SV1 on tumor growth and maintenance. Hormone ablation therapy is the primary clinical treatment for advanced prostate cancer. The ARR_2_PB promoter driving c-MYC expression contains two androgen response elements, which increases the level of c-MYC expression in an androgen-dependent manner, therefore, androgen ablation can reduce *c-MYC* expression. Since the ARR_2_PB promoter is regulated by androgens, surgical castration can serve as a method to induce the loss of *c-MYC* expression. However, the effects of androgen ablation must also be considered in this setting. Castrated (*n* = 5) (“c-MYC off”) and intact littermates (*n* = 5) (“c-MYC on”) were sacrificed 3 months after castration for analysis (Figure 4A). Gross and histological analysis of castrated and control mice were performed to evaluate the phenotypic effects of castration. MYC-SV1 urogenital systems post-castration weighed significantly less suggesting sustained c-MYC expression was required for MYC-SV1 tumor maintenance (*P* = 0.05) (Figure 4B). We examined the consequences of castration on the MYC-SV1 prostate. H&E revealed a significant reduction in prostate tumor lesions with the presence of involuted, atrophic, fibrotic lumens, increased deposition of collagen, with substitution by adipose tissue in castrated mice (Figure 4C-E). A decrease in c-MYC transgene expression by IHC was observed, lesions retained AR and KLF6-SV1 protein expression (Figure 4E-G). Cells within the castrated lumens expressed cytokeratin (CK) 18 (Figure 4H, brown). Dual staining with CK5 (green), a basal cell specific cytokeratin, confirmed the loss of basal cells in tumors of intact mice, a sign of invasion. The presence and restructuring of basal epithelial cells around luminal epithelial cells were noted in castrated mice (Figure 4H) (*P* = 0.0017). Staining for α-SMA identified the smooth muscle cells of the prostate (Figure 4I). In the intact mice, smooth muscle cells normally encircling prostate glands were displaced, however, these cells encircled the atrophic and mPIN glands of the castrated mice, reforming the natural prostate gland architecture (*P* = 0.0151). Cells remained highly proliferative as evidenced by strong nuclear PCNA expression (*P* = 0.5784) suggesting that KLF6-SV1 is sufficient to maintain mPIN and androgen-independent growth of the prostate (Figure 4J).

**Figure 4.**
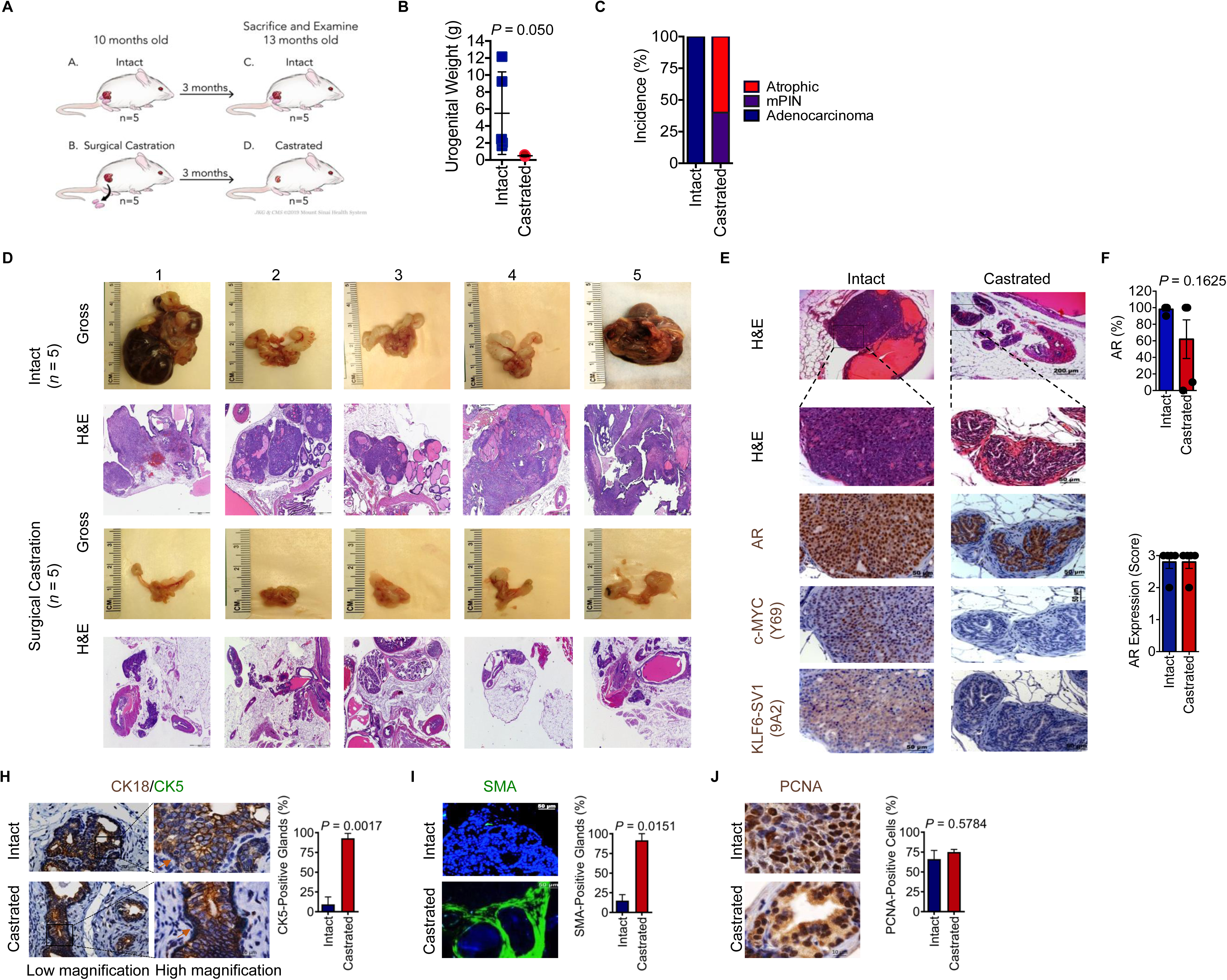
Castration and de-induction of c-MYC on the maintenance of MYC-SV1 prostate tumors. **A**, Schematic of castration experiment using the MYC-SV1 GEMM. Mice were castrated at 10 months of age (n = 5) post-tumor development and were sacrificed 3 months post-surgery. **B**, Urogenital weights of intact or castrated MYC-SV1 mice (mean ± s.d., *P* = 0.05, Student’s t-test). **C**, Incidence of mPIN and adenocarcinoma in intact and castrated mice. **D**, Gross images and micrographs of H&E of the urogenital system of all mice in the study. **E**, Histological evaluation of treatment phenotype incidence (%). H&E and immunohistochemical (IHC) staining of AR, c-MYC and KLF6-SV1 (10x magnification, scale bar=200µm; 40x magnification, scale bar=50µm). **F,** Percent of epithelial cells expressing AR (mean ± s.d., *P* = 0.1625, Student’s t-test). A total of more than 500 cells were counted from high-power fields. **G,** AR protein expression intensity. AR expression was measured on a 1+, 2+ or 3+ scoring system (0=negative, 1=weak, 2=moderate, 3=strong). Samples scored by intensity. Intensity distribution with representative images; primary tumors with total regions scored. **H,** IHC staining of CK5 (green) (mean ± s.d., *P* = 0.0017, Student’s t-test) and CK18 (brown) of intact and castrated prostate tissue (63x magnification, scale bar=20µm; 100x magnification, scale bar=10µm). **I,** Representative immunofluorescence staining image for alpha-smooth muscle actin (SMA) (mean ± s.d., *P* = 0.0151, Student’s t-test) (40x magnification, scale bar=50µm). **J,** Representative IHC staining image of PCNA (100x magnification, scale bar=10µm) with percent of epithelial cells expressing PCNA (mean ± s.d., *P* = 0.5784, Student’s t-test). A total of more than 500 cells were counted from high-power fields.

### Proteomic analysis reveals KLF6-SV1-promoting epithelial de-differentiation through the upregulation of vimentin

Having observed the progression to metastatic disease in the MYC-SV1 GEMM model, we sought to use unbiased proteomic profiling to better understand the pathway perturbations in our GEMM. Specifically, we used a proteome-wide profiling using label-free mass spectrometry (LC-MS/MS assays) of the WT, KLF6-SV1, Hi-Myc, and MYC-SV1 GEMMs (Figure 5A). Primary tissue was harvested at 10 months of age from each model for protein profiling; 10,677 peptides corresponding to 2,142 proteins (as determined by unique HUGO gene symbols) were analyzed, of which 1085 showed statistically significant differential expression (*P* < 0.01). Principal component analysis showed clear segregation of the samples by transgene expression (Figure 5B). A protein cluster was identified, “group B” (Figure 5C), that showed increased expression in the MYC-SV1 tumors compared to Hi-Myc or KLF6-SV1 prostates alone. Hierarchical clustering analysis revealed vimentin as a major phenotypic biomarker (Figure 5C, 5D). Furthermore, Ingenuity Pathway Analysis (IPA) also highlighted vimentin as among the top upregulated proteins in MYC-SV1 tumors (Figure S7). The convergence of hierarchical clustering and IPA analysis supported vimentin as playing a key role in the transformation of MYC-SV1 tumors. Unbiased proteomic data reinforced the finding that coordinate expression of KLF6-SV1 and c-MYC resulted in changes in prostate cancer cellular plasticity and dedifferentiation. Protein analysis by Western blot also confirmed increased vimentin protein expression in MYC-SV1 tumors compared to the prostates from the other GEMM mice analyzed (Figure 5E). Next, we examined vimentin expression by IHC to assess protein expression in individual cells while preserving the spatial tissue context of the WT, KLF6-SV1, Hi-Myc, and MYC-SV1 (Figure 5F). Vimentin was predominantly expressed in the connective tissue of WT and Hi-Myc mice.

**Figure 5.**
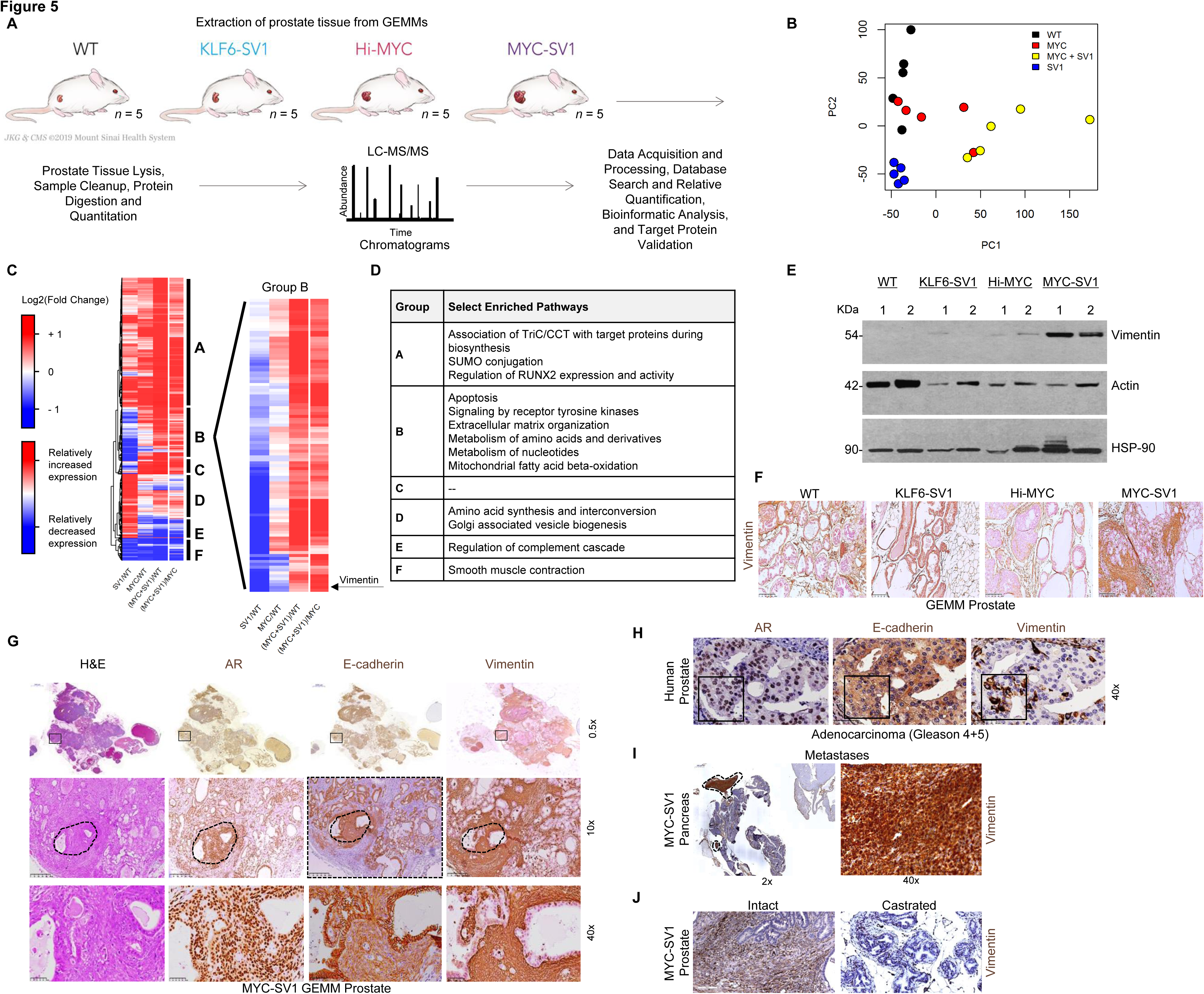
Proteomics analysis identifies a distinct expression signature with significant upregulation of vimentin in MYC-SV1 mouse model. **A**, Proteomics analysis experimental design. **B,** Principal component analysis (PCA) including WT (*n*=5; black dots), KLF6-SV1 (*n*=5; blue dots), Hi-Myc (*n*=5; red dots) and MYC-SV1 (*n*=5; yellow dots) prostate tissue specimens. **C**, Heatmap of significantly altered proteins amongst GEMM groups normalized to WT or Hi-Myc, as indicated (*P* < 0.01, ANOVA). A magnified subset of the heatmap (group B) is shown to the right. **D,** Select enriched pathways. **E,** Western blot of vimentin protein expression of mouse prostates. **F,** IHC staining of vimentin shows increased expression of vimentin in MYC-SV1 tumors (10x, scale bar=250µm). **E,** MYC-SV1 lesions expressing high levels of vimentin protein expression also concurrently express AR and E-cadherin confirmed by IHC staining of consecutive FFPE tumor sections of MYC-SV1 prostate (10x magnification, scale bar=250µm). Representative areas are noted with an asterisk (*) and shown at high magnification (40x magnification, scale bar=50µm). **H**, IHC staining of consecutive sections of human prostate cancer for vimentin expression (10x magnification, scale bar=250µm; 40x magnification, scale bar=50µm). **I,** IHC staining of MYC-SV1 pancreatic metastases for vimentin (encircled by dashed lines). High and low magnification photomicrographs are shown (2x magnification, scale bar=500µm; 40x magnification, scale bar=50µm). **J**, IHC staining of vimentin expression in intact vs. castrated MYC-SV1 mouse prostate (10x magnification, scale bar=250µm; 40x magnification, scale bar=50µm).

Cells expressing elevated vimentin protein were largely located within poorly differentiated tumor foci in the MYC-SV1 model (Figure 5F, G). Molecular characterization by IHC staining of vimentin, E-cadherin, and the AR, on consecutive tumor sections, revealed the co-expression of AR and E-cadherin in cells that expressed elevated vimentin (Figure 5G). These changes in protein expression were also observed in high-grade human prostate adenocarcinoma (Figure 5H). MYC-SV1 pancreatic metastases (see Figure 2) also expressed elevated vimentin (Figure 5I). Vimentin expression was absent in lesions of castrated MYC-SV1 mice (“c-MYC-off”) compared to lesions of intact mice (“c-MYC-on”) (see Figure 4) (Figure 5J). We examined the dual expression of vimentin and E-cadherin in Hi-Myc and MYC-SV1 mice by dual staining (Figure 6A). In well-differentiated lesions of Hi-Myc and MYC-SV1 GEMMs, expression of E-cadherin was localized to the cell membrane and vimentin expression was absent. Poorly differentiated lesions of MYC-SV1 predominately expressed vimentin with decreased and diffuse E-cadherin co-expression (Figure 6A). Western blot analysis confirmed an induction in vimentin protein expression with a concomitant decrease in E-cadherin protein expression in MYC-SV1 prostate tumors compared to those of Hi-Myc mice (Figure 6B). To confirm the role of KLF6-SV1 driving epithelial plasticity with characteristics of epithelial to mesenchymal transition, we turned to *in vitro* gene expression analysis. Exogenously expressed KLF6-SV1 in the Myc-CaP (KLF6-SV1-negative) mouse prostate epithelial cell line isolated from Hi-Myc mice (Ellwood-Yen K et al., 2003) led to a concomitant increase in c-MYC and vimentin protein expression with a decrease in E-cadherin protein expression (Figure 6C). We examined *CDH1* and *VIM* mRNA and E-cadherin and vimentin protein expression in a panel of human prostate cell lines of differing disease states (see Figure 3). *CDH1* mRNA and E-cadherin protein expression were significantly decreased in prostate cancer cell lines compared to non-tumorigenic cell lines (*P* = 0.0189, Figure 6D, 6F), while *VIM* mRNA and vimentin protein expression were significantly increased in prostate cancer cell lines compared to non-tumorigenic cell lines (*P* = 0.0029, Figure 6E, 6F). To further confirm that the phenotype observed was due to the cooperativity between c-MYC and KLF6-SV1, we examined the expression of vimentin and E-cadherin in the RWPE-1 stable non-tumorigenic cell lines and siRNA-treated PC3 cell lines shown in Figure 3. RWPE-1 cells stably expressing c-MYC and/or KLF6-SV1 also demonstrated a concomitant decrease in *CDH1* mRNA and E-cadherin protein expression (*P* < 0.0001, Figure 6G, 6I) with a coordinate upregulation of vimentin mRNA and protein expression (*P* < 0.0001, Figure 6H, 6I). Collectively, these cellular and *in vivo* data suggest that KLF6-SV1, in the presence of c-MYC, regulated vimentin and E-cadherin expression, inducing cellular plasticity promoting prostate cancer progression and metastasis (Figure 6J).

**Figure 6.**
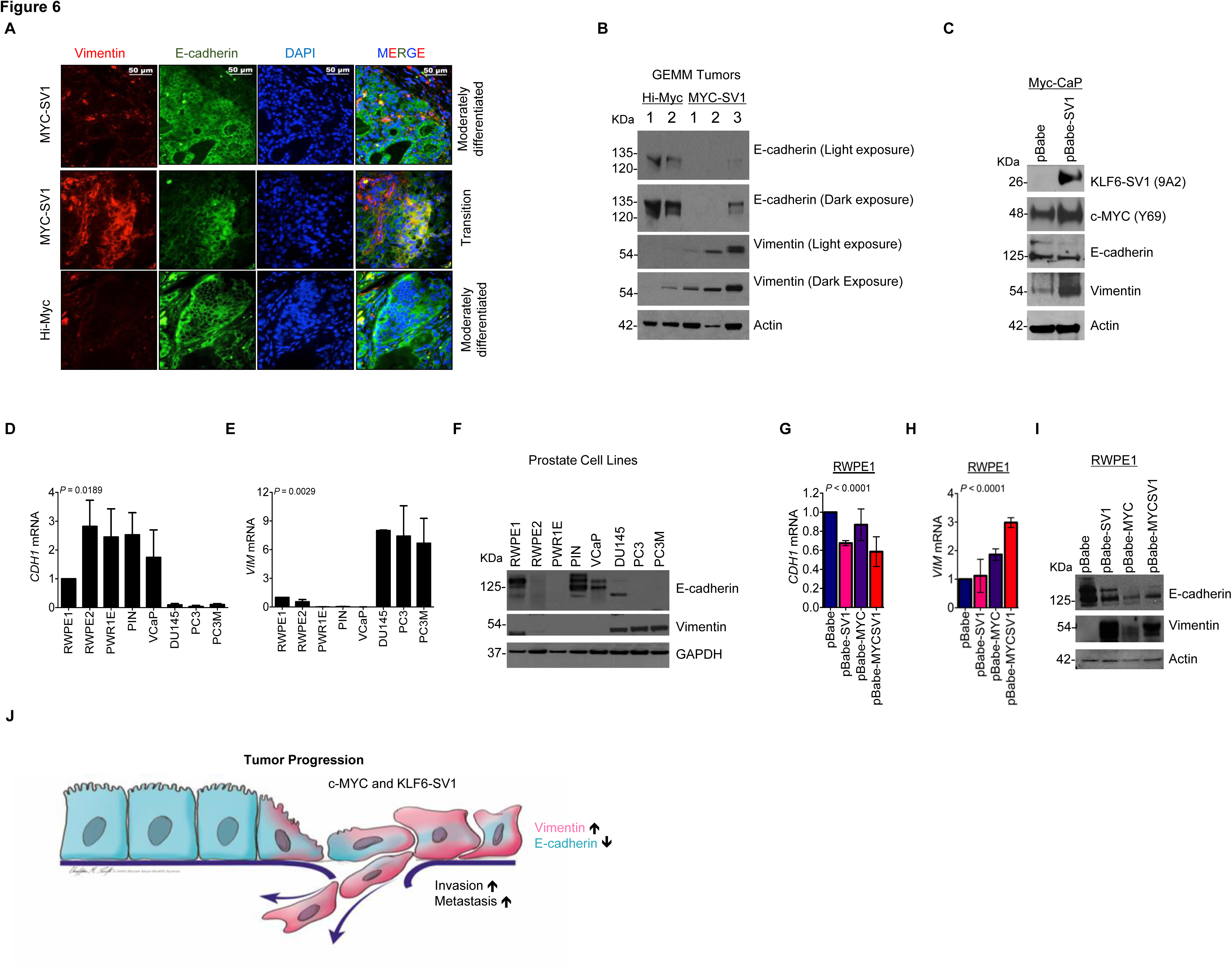
KLF6-SV1 dependent up-regulation of vimentin. **A**, Micrographs of immunoflourescence (IF) staining for vimentin (red) and E-cadherin (green) capturing single and dual-positive tumor cells (yellow). **B**, Western blot analysis of vimentin protein expression in MYC-SV1 and Hi-MYC prostate tumors. **C**, Western blot of exogenous expression of KLF6-SV1 in Myc-CaP cells for c-MYC, KLF6-SV1, vimentin and E-cadherin protein expression. **D**, E-cadherin mRNA in a panel of prostate cell lines of differing disease states (mean ± s.d., *P* = 0.0189, ANOVA). Fold change to RWPE-1 expression. Bars represent means ± s.d. of three biological replicates. **E**, qRT-PCR for vimentin mRNA expression in a panel of prostate cell lines of differing disease states (mean ± s.d., *P* = 0.0029, ANOVA). Fold change normalized to RWPE-1 expression. Error bars represent s.d. and experiment performed in triplicate. **F**, E-cadherin and vimentin protein expression in a panel of prostate cell lines of differing disease states. **G**, qRT-PCR for E-cadherin mRNA expression of stable cell lines over-expressing p-Babe-KLF6-SV1, p-Babe-c-MYC, or p-Babe-KLF6-SV1/c-MYC in RWPE-1 cells (mean ± s.d., *P* < 0.0001, ANOVA). Bars represent means of three biological replicates. **H**, qRT-PCR of vimentin mRNA expression of stable cell lines over-expressing p-Babe-KLF6-SV1, p-Babe-c-MYC, or p-Babe-KLF6-SV1/c-MYC in RWPE-1 cells (*P* < 0.0001). Bars represent means of three biological replicates. **I**, E-cadherin and vimentin protein expression in stable RWPE-1 cell lines over-expressing p-Babe-KLF6-SV1, p-Babe-c-MYC or p-Babe-KLF6-SV1/c-MYC. Bars represent means of three biological replicates. **J**, Schematic of KLF6-SV1/c-MYC-induced dedifferentiation.

We next sought to determine whether concurrent c-MYC and KLF6-SV1 expression occurred in human prostate cancer by RNA *in situ* hybridization (RNA ISH) and IHC staining. In our proof-of-concept study of human prostate cancer and liver metastases from 19 patients (57 specimens) (Figure 7A), c-MYC and KLF6-SV1 expression were significantly positively correlated at the RNA (Figure 7B, 7C) and protein levels (Figure 7D, 7E). Increased epithelial c-MYC mRNA as evaluated by *in situ* hybridization was significantly associated with increased KLF6-SV1 mRNA expression (Pearson *r* = 0.7301, *P* < 0.0001 and Spearman *r* = 0.8041*, P* < 0.0001). c-MYC protein expression assessed by IHC was associated with significantly increased KLF6-SV1 protein (Pearson *r* = 0.5424, *P* < 0.0001 and Spearman *r* = 0.4983, *P* < 0.0001). These findings provide a proof-of-principle of our working hypothesis of the cooperation between c-MYC and KLF6-SV1 in human prostate tumorigenesis.

**Figure 7.**
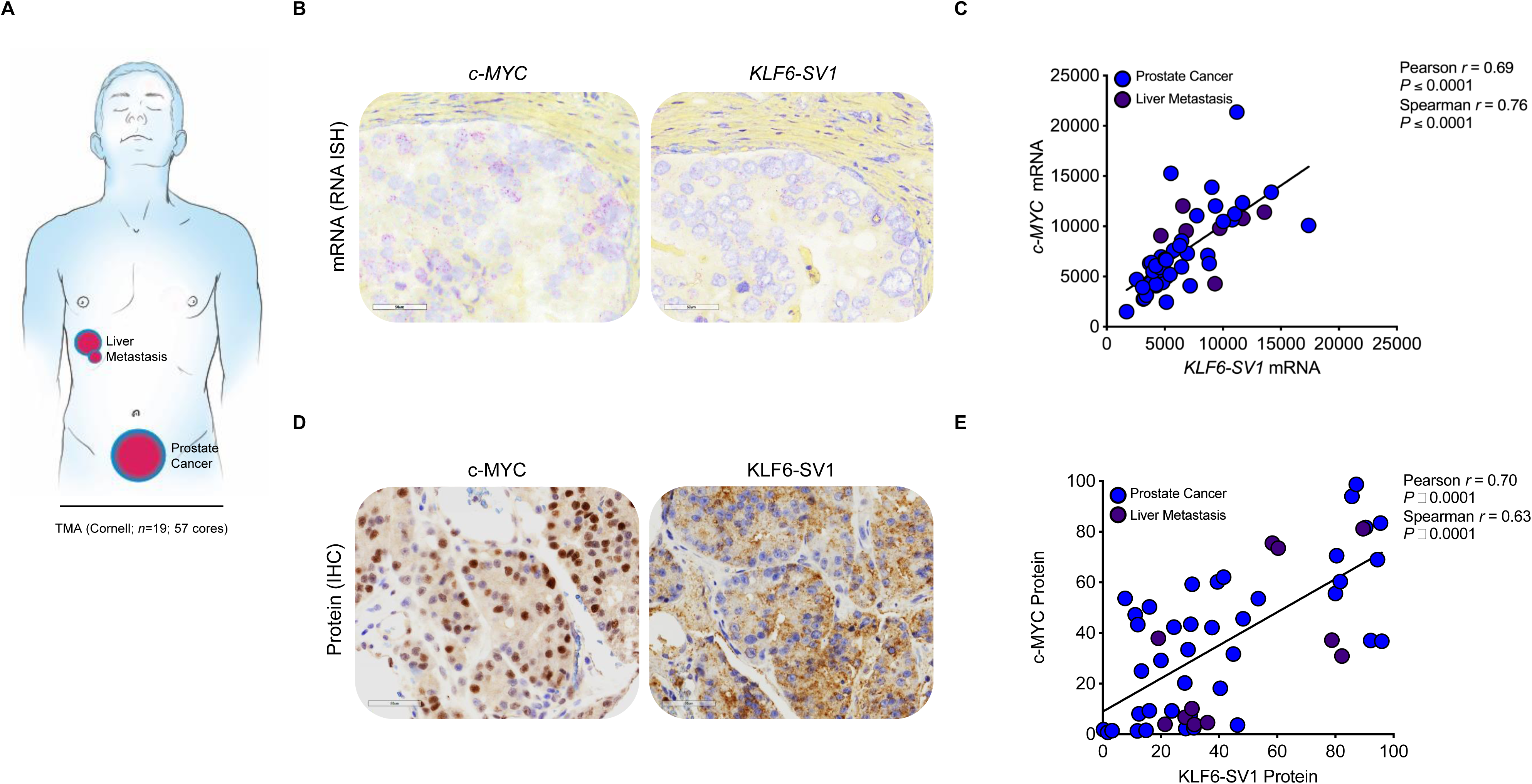
KLF6-SV1 and c-MYC are correlated events in the human prostate. **A**, Schematic of primary (*n* = 48 cores; 16 patients) and metastatic (liver) (*n* = 9 cores; 3 patients) specimens included in TMA for c-MYC and KLF6-SV1. **B**, Representative RNA ISH staining with specific probes against c-MYC or KLF6-SV1. Signals are granular and discrete red dots corresponding to individual RNA targets. **C,** Quantification and correlation between *c-MYC* and *KLF6-SV1* mRNA (Pearson *r* = 0.59, *P* < 0.0001 and Spearman *r* = 0.71, *P* < 0.0001). **D**, Representative immunohistochemical staining with specific antibodies against c-MYC or KLF6-SV1. **E**, Quantification and correlation between c-MYC and KLF6-SV1 protein observed in human prostate specimens (Pearson *r* = 0.70, *P* < 0.0001 and Spearman *r* = 0.63, *P* < 0.0001).

## Discussion

The goal of this study was to investigate oncogenic events that govern the genesis and progression of aggressive and metastatic prostate cancer. Defining the discrete genetic events that lead to prostate cancer development and metastases are critical for generating preclinical models to study the lethal form of prostate cancer. The molecular basis of prostate cancer has been explored by several groups revealing numerous molecular alterations with the potential to act as drivers of cancer development and progression (Dhanasekaran SM et al., 2001, Lapointe J et al., 2004, Taylor BS et al., 2010, Grasso CS et al., 2012). The functional and biological impact of some of these genetic alterations has been defined through the generation and characterization of GEMMs. One identified dysregulated gene, *c-MYC*, is recognized as a major contributor to the onset of mPIN, a key precursor lesion to adenocarcinoma, and prostate cancer initiation and maintenance, in both mouse and human (Zhang X et al., 2000, Ellwood-Yen K et al., 2003, Ellis L et al., 2016, Nguyen HG et al., 2018, Watson PA et al., 2005). Elevated *c-MYC* mRNA and protein expression is present in greater than 50% of metastatic prostate cancer (Taylor BS et al., 2010, Beltran H et al., 2016), thus, raising questions about the additional genetic events that cooperate with *c-MYC* to induce the metastatic cascade. We demonstrate that cooperativity between two frequent genetic events in human prostate cancer, increased expression of *c-MYC* and *KLF6-SV1*, are necessary and sufficient to promote and drive progression to a poorly differentiated and metastatic state. Our results have uncovered a mechanism underlying prostate cancer progression and demonstrates that *c-MYC* and *KLF6-SV1* cooperate to accelerate prostate tumor development from localized to metastatic disease and induce cellular plasticity. In benign prostate glands, *KLF6-SV1* expression induces pre-malignant transformation (i.e., mPIN). In the presence of c-MYC, KLF6-SV1 profoundly alters c-MYC-driven prostate tumorigenesis at an early stage, resulting in a more rapid onset of cancer, leading to poorly differentiated, invasive, and metastatic disease with a unique proteomic program, including significant upregulation of vimentin in MYC-SV1 prostate tumors. Taken together, our findings implicate KLF6-SV1 as a key regulator of cellular plasticity, invasion, and metastasis. We found that KLF6-SV1 was necessary and sufficient to drive the plasticity of tumor cells in the presence of c-MYC. A predominant subset of these tumor cells displayed markedly elevated vimentin expression with concurrent loss of E-cadherin, characteristic molecular features of epithelial to mesenchymal transition, invasion, and metastasis. Defects in epithelial cell-cell adhesion and cell polarity have been associated with prostate cancer initiation, progression, and aggressiveness (Das R et al., 2014). Dedifferentiation and cellular plasticity have been described in many tumor types, including prostate cancer, as a contributor of tumor heterogeneity, metastasis, treatment resistance, disease recurrence, and poor outcomes (Das R et al., 2014, Yang J et al., 2020). Our histological and molecular findings corroborate with previously reported clinical outcomes in patients with advanced and metastatic prostate cancer (Zhao Y et al., 2008, Armstrong AJ et al., 2011, Lang SH et al., 2002). Lang et al. and Zhao et al. found that vimentin expression to be clinically relevant as it significantly correlated with prostate cancer disease aggressiveness. High vimentin protein expression was observed in tumors of men with poorly differentiated and metastatic prostate cancer to the bone compared to well– and moderately differentiated tumors (Lang SH et al., 2002, Hanahan D and Weinberg R, 2011). Additionally, the expression of E-cadherin was demonstrated to be inversely associated with vimentin expression in human prostate cancer patients (Armstrong AJ et al., 2011). Circulating tumor cells isolated from patients with metastatic prostate cancer expressed vimentin and E-cadherin (Armstrong AJ et al., 2011).

Our loss– and gain-of-function studies confirmed the mutually regulated nature of this cooperativity and that of KLF6-SV1 expression is responsible for the significant changes in E-cadherin and vimentin protein expression observed. Furthermore, inactivation of an initiating oncogene suggests that a single oncogene can play a crucial role in the maintenance of genetically complex tumors (Podsypanina K et al., 2008, Soucek L et al., 2008). In our model system, androgen deprivation reflected the loss of c-MYC oncogene expression (Ellwood-Yen et al., 2003). Our findings demonstrate that interfering with cooperating genetic events can induce regression, a finding that may have important implications regarding prostate cancer treatment. In the absence of *c-MYC*, *KLF6-SV1* was insufficient to drive full and rapid malignant transformation. c-MYC played a crucial role in the maintenance and viability of MYC-SV1 tumors, while KLF6-SV1 acted as a potent driver of MYC-SV1 tumor progression and aggressiveness in the presence of c-MYC. Additionally, prostates from castrated mice retained expression of AR and PCNA suggesting that KLF6-SV1 signaling may play a role in androgen-independent growth of prostate epithelial cells, a characteristic not observed in castrated Hi-Myc mice (Ellwood-Yen K et al., 2003). A longer treatment course is needed to determine if these mice ultimately develop castration resistant prostate cancer. Collectively, our findings are consistent with other c-MYC-driven tumor mouse models (e.g., sarcoma and liver cancer) and those displaying cooperation with RAS (e.g., breast cancer and lung cancer) in which the inactivation of c-MYC expression led to the regression of tumors demonstrating the necessity for c-MYC in tumor maintenance (Podspanina K et al., 2008, Soucek L et al., 2008, Shachaf CM et al., 2004, Jain M et al., 2002).

The human relevance of our MYC-SV1 GEMM is credentialed by the significant correlation of these genetic alterations in human tumors, and the recapitulation of clinically relevant phases of human prostate cancer progression histologically. MYC-SV1 tumors also possess characteristics that define the “hallmarks of cancer”: sustaining proliferation, evading growth suppressors and cell death, limitless replicative ability, angiogenesis, and invasion and metastasis (Hanahan D and Weinberg R, 2011). In addition to our genetic and experimental evidence in the GEMMs, corroborating results were obtained in prostate cell lines of various histological origins and validated by the significant correlation of these genetic alterations in humans. The novelty of our work resides in unraveling the genetic cooperativity between *c-MYC* and *KLF6-SV1* revealing primary and metastatic prostate cancer development at a rate greater than the additive effect of each oncogenic factor alone. The enhanced and reliable tumor kinetics have implications in the multistep nature of tumor progression and metastasis and contribute to our understanding of the genetic drivers of lethal disease. In summary, our findings further improve our understanding of the genetic drivers of lethal prostate cancer and identify promising targets for biomarker and therapeutic development.

## Methods

### Cell Lines

Prostate (RWPE-1, RWPE-2, PWR1E, VCaP, DU145, PC3, PC3M) and phoenix cell lines were obtained from the American Type Culture Collection (Manassas, VA). Myc-CaP and Prostatic Intraepithelial Neoplasia (PIN) cell lines were a generous gift from Dr. Charles Sawyers (Memorial Sloan Kettering, New York, NY) and Dr. Mark Stearns (Drexel University, Philadelphia, PA) respectively. All cells were maintained at 37°C with 5% CO_2_. RWPE-1, RWPE-2, PWR-1E and PIN cells were cultured in Keratinocyte Serum-Free Medium supplemented with bovine pituitary extract and recombinant human epidermal growth factor (Gibco, Life Technologies, Grand Island, NY). Myc-CaP, VCaP, DU145, PC3, and PC3M cells were cultured in Dulbecco’s modified Eagle’s medium (HyClone, GE Healthcare Life Sciences, Logan, UT) supplemented with 10% fetal bovine serum (HyClone, GE Healthcare Life Sciences, Logan, UT) and 0.5% penicillin-streptomycin (Life Technologies, Grand Island, NY). Cells were regularly tested for mycoplasma with MycoAlert per manufacturer’s protocol (LT07-318, Lonza, Basel, Switzerland).

### Exogenous expression or siRNA knockdown

Stable cell lines were generated by retroviral infection with pBABE-puro (1764, Addgene, Cambridge, MA), pBABE-zeo (1766, Addgene, Cambridge, MA), pBabe-c-myc-zeo (17758, Addgene, Cambridge, MA) or pBabe-KLF6-SV1 virus. Retrovirus was generated by transfecting oncogenic plasmids and viral packaging plasmid (pCL-Ampho) into phoenix cells with Lipofectamine 2000 (Life Technologies, Grand Island, NY). Viral particles were incubated with prostate cells (70% confluence) in polybrene (4 mg/ml) for 24 hours. Infected cells were selected for puromycin or zeocin resistance (2 mg/mL). Targeted knockdown was conducted with 100nM ON-TARGETplus smartpool human *c-MYC* (mixture of 4 individual siRNA) and/or human *KLF6-SV1* siRNA (Dharmacon, Lafayette, CO) with HiPerfect (Qiagen, Valencia, CA).

### Quantitative real-time PCR

RNA was extracted using the RNeasy Mini Kit according to the manufacturer’s protocol and treated with DNase (Qiagen, Valencia, CA USA). cDNA was prepared with the Fermentas cDNA synthesis kit according to the manufacturer’s instructions. mRNA levels were quantified by quantitative real-time polymerase chain reaction (qRT-PCR) using the following PCR primers on a ABI PRISM 7900HT Sequence Detection System (Applied Biosystems). Primer sequences: KLF6-SV1 endogenous F: 5′-CCTCGCCAGGGAAGGAGAA-3′, R: 5′-CGGTGTGCTTTCGGAAGTG-3′; KLF6-SV1 exogenous F: 5′-CCTCGCCAGGGAAGGAGAA-3′, R: 5′-AAAACGCCACTCACACC-3′; c-MYC exogenous F: 5′-TTTCGGGTAGTGGAAAACCA-3′, c-MYC R: 5′-GAGGAGGAGCAGCGTCATCT-3′; c-MYC endogenous F: 5′-TTCGGGTAGTGGAAAACCAG-3′, R: 5’ CAGCAGCTCGAATTTCTTCC-3′; KI-67 F: 5’-ATCGTCCCAGGTGGAAGAGTT-3’, R: 5’-ATAGTAACCAGGCGTCTCGTGG-3’; E-cadherin (CDH1) F: 5′-CAAAGTGGGCACAGATGGTGTG-3′, R: 5′-CTGCTTGGATTCCAGAAACGG-3; Vimentin (VIM) F: 5′-CAGATTCAGGAACAGCATGTC-3′, 5′-TCAGAGAGGTCAGCAAACTTG-3′. Quantitative real-time PCR was performed on an ABI PRISM 7900HT Fast Real-Time PCR machine using SYBR Green (Applied Biosystems, Foster City, CA). Data was analyzed with SDS 2.3 software. Expression levels were determined using the ΔCt method normalized to GAPDH, β-Actin and 18S. Graphs shown are of expression levels normalized to GAPDH.

### Western blot

Tissues and cell protein lysates were prepared with RIPA buffer (89900, Thermo Fisher Scientific, Waltham, MA), which includes a protease inhibitor cocktail (05892791001, Roche/Sigma-Aldrich, St. Louis, MO). Proteins were quantified by the protein quantification assay (BioRad, Hercules, CA). Equal amounts of protein (40-50µg) were denatured and separated on 12% Bis-Tris SDS–polyacrylamide electrophoresis gels (NP0341BOX, Invitrogen, Carlsbad, CA) and transferred to a nitrocellulose membrane (BioRad, Hercules, CA). Membranes were blocked with 5% nonfat milk (LabScientific Inc., Livingston, NJ) in Tris-Buffered Saline–Tween (77500, Affymetrix, Thermo Fisher Scientific, Waltham, MA) and incubated, at 4°C overnight with antibodies against KLF6 (sc-7158, Santa Cruz Biotechnology Inc., Dallas, TX), KLF6-SV1 (9A2, Hybridoma Facility, Icahn School of Medicine at Mount Sinai), E-cadherin (sc-7870, Santa Cruz Biotechnology Inc., Dallas, Texas, USA; ab40772, Abcam, Cambridge, MA), Vimentin (ab92547, Abcam, Cambridge, MA, USA; sc-6260, Santa Cruz Biotechnology Inc., Dallas, TX; Epitomics 2707-1, Burlingame, CA 94010), c-MYC (ab32072, clone Y69, Abcam, Cambridge, MA; OP10, clone 9E10, Calbiochem, San Diego, CA), GAPDH (8884, Cell Signaling, Danvers, MA; sc-25778, Santa Cruz Biotechnology Inc., Dallas, TX), Actin (12620, Cell Signaling, Danvers, MA; sc-1616, Santa Cruz Biotechnology Inc., Dallas, TX), HSP-90 (79641S, Cell Signaling, Danvers, MA) anti-mouse (31430, Thermo Fisher Scientific, Rockford, IL) or anti-rabbit (Jackson ImmunoResearch Laboratories, West Grove, PA) horseradish peroxidase–conjugated secondary antibody. Membranes were developed using enhanced chemiluminescence (Lumi Light or Lumi Light-plus, Roche/Sigma-Aldrich, St. Louis, MO) following the manufacturer’s instructions.

### Thymidine proliferation assay

Proliferation was determined by estimating [3H] thymidine incorporation. 1 μCi/ml [3H] thymidine (Amersham, Arlington Heights, IL) was added to cells. After 2-hr incubation period, cells were washed with ice-cold PBS and fixed in methanol at 4°C. Cells were solubilized in 0.25% sodium hydroxide and 0.25% Sodium Dodecyl Sulfate, Molecular Biology Grade (SDS) and neutralization with hydrochloric acid (1N). Disintegrations per minute were estimated by liquid scintillation counting.

### Cell cycle analysis

Cells were harvested, fixed in absolute ethanol (Sigma-Aldrich, St. Louis, MO), stained with a solution of propidium iodide, RNase A, and PBS, and analyzed by the FACScan flow cytometer. The data was analyzed with CellQuest software (BD Biosciences, San Jose, CA).

### Establishment of mouse colonies and genotyping

All mice were housed in the animal facility at the Icahn School of Medicine at Mount Sinai. Mice were cared for in accordance with institutional guidelines to ensure humane, responsible, and appropriate care of the animals (IACUC #08-0034). Mice were maintained on a 12-hour light/dark cycle, at a constant temperature and relative humidity. Tap water and food were available *ad libitum*. Hi-Myc mice were obtained from the Mouse Repository of the National Cancer Institute Mouse Models of Human Cancer Consortium at National Cancer Institute (Frederick, MD). KLF6-SV1 mice were kindly provided by Dr. John Martignetti (Icahn School of Medicine at Mount Sinai). KLF6-SV1 mice were engineered to express human KLF6-SV1, which was cloned into the pCAGG-neo expression vector that is driven by the β-actin promoter. Following pronuclear injection of the purified construct, a total of fourteen pups were produced and analyzed. Three of six pups (s1–s3) in the first of two litters contained the KLF6-SV1 insert as revealed by quantitative real-time PCR (qRT-PCR) of the human-specific 602-bp sequence in the KLF6-SV1 gene. Three independent KLF6-SV1 transgenic founders were generated. The Mouse Genetics and Gene Targeting Core at the Icahn School of Medicine at Mount Sinai performed injections into FVB blastocysts. KLF6-SV1 transgenic mice were crossed with Hi-Myc mice and germline transmission was determined by genotyping. Mouse tail DNA was isolated and genotyped to evaluate transgene transmission with Phire Animal Tissue Direct PCR (F-140, Thermo Fisher Scientific, Waltham, MA).

### Histopathological assessment and tissue analysis

Mouse prostates were isolated and fixed in 10% neutral-buffered formalin (SF100-4, Thermo Fisher Scientific, Waltham, MA) overnight, paraffin-embedded, sectioned and stained with haematoxylin and eosin (H&E). Murine model characterization and documentation of lesion progression was conducted in a blinded manner following the Bar Harbor Classification System (Shappell SB et al., 2004). All mice were age-matched and of identical genetic background (FVB/N inbred background). For immunohistochemistry, 5µm sections were deparaffinized with xylene and rehydrated through graded ethanol washes followed by antigen retrieval in sodium citrate buffer (10 mM, pH 6.0) with a pressure cooker (Dako, Carpinteria, CA), blocked, and incubated with primary antibodies AR (ab133273, Abcam, Cambridge, MA, USA), vimentin (ab92547, Abcam, Cambridge, MA; 9855S (Alexa Fluor 555), Cell Signaling, Danvers, MA), E-cadherin (ab40772, Abcam, Cambridge, MA; 3199S (Alexa Fluor 488), Cell Signaling, Danvers, MA), Smooth Muscle Actin (ab188498, Abcam, Cambridge, MA), c-MYC (ab32072, Abcam, Cambridge, MA), PCNA (ab92552, Abcam, Cambridge, MA), KLF6-SV1 (9A2, Hybridoma Facility, Icahn School of Medicine at Mount Sinai), or NKX3.1 (AB5983, MilliporeSigma, Burlington, MA) overnight at 4°C. Sections were washed with PBS (PBS940M, Biocare Medical, Concord, CA). The Mouse on Mouse (MOM) Detection Kit (BMK-2202, Vector Laboratories, Burlingame, CA) was used for mouse primary antibodies. EnVision+ Rabbit (K400311-2, Agilent Technologies, Inc., Santa Clara, CA) was used as a secondary antibody. Proteins were visualized with 3,3′-diaminobenzidine (DAB) (K346811-2, Agilent Technologies, Inc., Santa Clara, CA) or Vina Green (BRR807AS, Biocare Medical, Concord, CA). Sections were counterstained with hematoxylin (CATHE-H, Biocare Medical, Concord, CA), dehydrated, and mounted with Permount (Thermo Fisher Scientific, Waltham, MA). For immunofluorescence staining, tissue was counterstained with DAPI nuclear stain (H-1200, Vector Laboratories, Burlingame, CA). Bright-field or fluorescent images were captured with Zeiss AxioImager Z2 (Jena, Germany). Images were captured at the Icahn School of Medicine at Mount Sinai Microscopy Core (supported by NIH P30CA196521). TUNEL staining was performed using the ApopTag Peroxidase *In Situ* Apoptosis Detection Kit (S7100, EMD Milipore, Billerica, MA) as per manufacturer protocol. For cell quantification, representative sections from more than three mice were counted for each genotype by using the cell counter function in ImageJ software (https://imagej.nih.gov/ij/).

## Castration treatment

MYC-SV1 mice were castrated at 10 months of age with definite adenocarcinoma (*n* = 5) and examined 3 months post-castration. The control group (intact) contained age-matched littermate animals (*n* = 5). Mice were anesthetized using isoflurane (Baxter Healthcare Corporation, Deerfield, IL). The perineal region was cleaned with ethanol and a 4–5 mm incision was made with sterile dissecting shears. Using sterile forceps, the testes were located, and a ligature was made around the testicular vessels and the tunica albuginea. The testes were amputated, and the scrotum was sutured. An antibiotic was applied over the region of the wound to help the healing process.

## Proteomics Analysis

### Sample preparation of WT, KLF6-SV1, Hi-Myc, and MYC-SV1 tissues for label-free expression

Tissues were lysed in 3% sodium dodecyl phosphate (SDS) and protease inhibitor (Sigma, part number 4693159001) using a pestle followed by pulse sonification. Once lysed, tissues were cleaned of detergent using a previously published filter-aided sample preparation protocol with a 10-kDa molecular weight cutoff filter (Millipore, Billerica, MA) and buffer exchanged with 8M Urea in 50mM Tris-pH-8.0 to a final volume of 50µL (Tomechko SE et al., 2015). After cleanup, protein concentration of each lysed sample was measured using the Bradford assay and following the manufacturer’s standard protocol (Bio-Rad, Hercules, CA). Next, 10μg of total protein were digested with Lysyl Endopeptidase (Wako Chemicals, Richmond, VA) at an enzyme:substrate ratio of 1:20 for 2-hours at 37°C. The urea concentration was then adjusted to 2M using 50mM Tris, pH 8, followed by an overnight trypsin digestion using sequencing grade trypsin (Promega, Madison, WI) at an enzyme:substrate ratio of 1:20 at 37°C. Digested peptides were then stored at −80°C until used for the downstream LC-MS/MS analysis.

### Reverse Phase LC-MS/MS Analysis

Six hundred nanograms of each sample were analyzed by LC-MS/MS using a LTQ-Orbitrap Elite mass spectrometer (Thermo Fisher Scientific, San Jose, CA) equipped with a nanoAcquity^TM^ Ultra-high pressure liquid chromatography system (Waters, Taunton, MA). The injection order on the LC-MS was randomized over all samples. Blank injections were run after each sample to minimize carry-over between samples. Mobile phases were organic phase A (0.1% formic acid in water) and aqueous phase B (0.1% formic acid in acetonitrile). Peptides were loaded onto a nanoACQUITY UPLC^®^ 2G-V/M C18 desalting trap column (180 μm x 20 mm nano column, 5μm, 100 Å) at flow rate of 0.300µl/minute. The analytical column used to resolve peptides was a nanoACQUITY UPLC^®^ BEH300 C18 reversed phase column (75μm x 250 mm nano column, 1.7μm, 100Å; Waters, Milford, MA). The gradient employed was 1-90 % of phase B over 240 minutes (isocratic at 1% B, 0-1 min; 2-42% B, 2-212 min; 42-90% B, 212-223 min; and 90-1% B, 223-240 min) at a flow rate of 300nL/min. A nano ES ion source at 1.5 kV spray voltage, and 270 °C capillary temperature was utilized to ionize peptides. Full scan MS spectra (m/z 380-1800) were acquired at a resolution of 60,000 followed by twenty data dependent MS/MS scans. MS/MS spectra were generated by collision-induced dissociation of the peptide ions (normalized collision energy = 35%; activation Q = 0.250; activation time = 20 ms) to generate a series of b-and y-ions as major fragments. LC-MS/MS raw data were acquired using the Xcalibur software (Thermo Fisher Scientific, version 2.2 SP1). LC-MS/MS raw data were then acquired using the Xcalibur software (Thermo Fisher Scientific, version 2.2 SP1).

### Data Processing for Protein Identification and Quantification

The LC-MS/MS raw files (one for each sample) were imported into Rosetta Elucidator™ (Rosetta Bio-software, version 3.3.0.1.SP.25) and processed as previously described. The peak list (.dta) files were searched by Mascot (version 2. 1, Matrix Science London, UK) against the Uniprot database. Mascot search settings were as follow: trypsin enzyme specificity; mass accuracy window for precursor ion, 10 ppm; mass accuracy window for fragment ions, 0.8 Da; carbamidomethylation of cysteines as fixed modifications; oxidation of methionine as variable modification; and one missed cleavage. Peptide identification criteria were a mass accuracy of ≤10 ppm, an expectation value of *P* < 0.05, and an estimated False Discovery Rate (FDR) of less than 2%. The search results were imported back into Elucidator and automated differential quantification of peptides was then accomplished with Rosetta Elucidator (Wisiewski JR et al., 2009). Normalization of signal intensities across samples was performed using the average signal intensities obtained in each sample.

### Bioinformatics analyses

Computational analyses of the processed dataset were performed with R version 3.3.2 unless otherwise noted. Differential peptide expression was evaluated by one-way ANOVA across all groups (WT, KLF6-SV1, Hi-Myc, and MYC-SV1). Principal component analysis (PCA) was calculated using the “prcomp” function and plotted using the “scatterplot3d” package. The heatmap of the protein intensities were generated with the “heatmap.2” function found in the “gplots” package. Each protein was represented by the peptide with the lowest one-way ANOVA statistic, and only proteins having a corresponding peptide with *P* < 0.01 were included in the heatmap. Those proteins were subsequently clustered using hierarchical clustering (“hclust”). The resulting dendogram was then divided into 6 groups using the “cutree” function. Proteins within each group were then analyzed for pathway enrichment by Fisher’s exact test using the PANTHER Statistical overrepresentation tool, released 20190711 (Mi H et al., 2019). The Reactome version 65 database of pathways was used as reference. The *P* values were corrected for multiple hypothesis testing using the Benjamini-Hochberg FDR method. Pathway analysis was performed using Ingenuity Pathway Analysis (Qiagen, Redwood City, CA; www.ingenuity.com) on protein, which had peptides that were deemed significant (*P* = 0.005 and a fold change > +/− of 3). Proteins identifications that met the above criteria for significance were imported into Ingenuity Pathways Analysis. Ingenuity Pathways Analysis calculated significant pathways using a right-tailed Fisher’s exact test. Top canonical pathways were chosen based on passing the significance criteria (*P* ≤ 0.05).

### *RNA ISH*, IHC, and analysis of human tissues

The tissue microarray (TMA) was constructed at Weill Cornell Medicine/New York Presbyterian Hospital (New York, NY). Archival pathology specimens from Weill Cornell Medicine/New York Presbyterian Hospital were obtained retrospectively under approved Institutional Review Board (IRB) protocol (IRB #1007011157). This study was also approved by the IRB at the Icahn School of Medicine at Mount Sinai (IRB#14-00757). The TMA contained a total of 57 prostate cancer specimens from 19 patients with localized (n=16 patients, 48 specimens) and metastases (n=3 patients, 9 specimens) and additional benign control tissues (prostate, liver, placenta, brain, kidney, and testis). H&E stained slides were reviewed by study pathologist (J.M.M.). RNA ISH was performed by the single-color chromogenic QuantiGene® ViewRNA ISH Tissue Assay (Affymetrix, Santa Clara, CA) using pairs of specially designed oligonucleotide probes that through sequence-specific hybridization recognizes both the specific target RNA sequence and the signal amplification system. A ViewRNA Type 1 target probe (1 bDNA) was designed for KLF6-SV1 (Affymetrix, Santa Clara, CA). ViewRNA Type 1 catalog probe was used for c-MYC (Affymetrix, Santa Clara, CA). Cross-hybridization to other sequences was minimized by screening against the entire human RNA sequence database. Signal amplification occurred at target sites bound by probe pairs only. Nonspecific off-target binding by single probes did not result in signal amplification. Quantigene ViewRNA ISH Tissue Control Kit (1-Plex), which includes GAPD, ACTB, PP1, E. Coli, K12 dapB, UBC, and PPIB (Affymetrix, Santa Clara, CA) was used as a staining technical control. Immunohistochemical (IHC) staining for c-MYC and KLF6-SV1 was applied using a commercially available antibody for c-MYC protein expression (monoclonal antibody, clone Y69, Abcam), and for KLF6-SV1 (monoclonal antibody, clone 9A2, Icahn School of Medicine at Mount Sinai) on the Leica Bond III automated stainer (Leica Biosystems Inc., Buffalo Grove, IL). Hybridization signals (red colorimetric staining) were detected under a brightfield microscope followed by counterstaining with hematoxylin. Signals were granular and discrete red dots corresponding to individual RNA targets. HALO™ Software (Indica Labs, Inc., Corrales, NM) was utilized for image analysis of RNA *in situ* expression and IHC staining for protein expression at a single cell resolution to quantify expression of positive epithelial cells (Park K et al., 2010).

## Materials Availability Statement

Materials of this study are available upon request from the corresponding author.

## Acknowledgments

The authors would like to acknowledge the support of the Howard Hughes Medical Institute (G.N.), NIH/NCI R01CA181654 (G.N., M.D.G., A.C.L., S.I.), NIH/OD U54OD020353 (C.C.C., S.I.), NIH/NCATS Loan Repayment Program (S.I.) and NIH/NCATS KL2RR029885 (S.I.). We thank Mireia Castillo-Martin, Michael Parides, and Raheleh Hatami for helpful discussions, and Jill Gregory and Christopher Smith of the Instructional Technology Group at the Icahn School of Medicine for illustrations.

## Supplemental Figures

**Figure S1.**
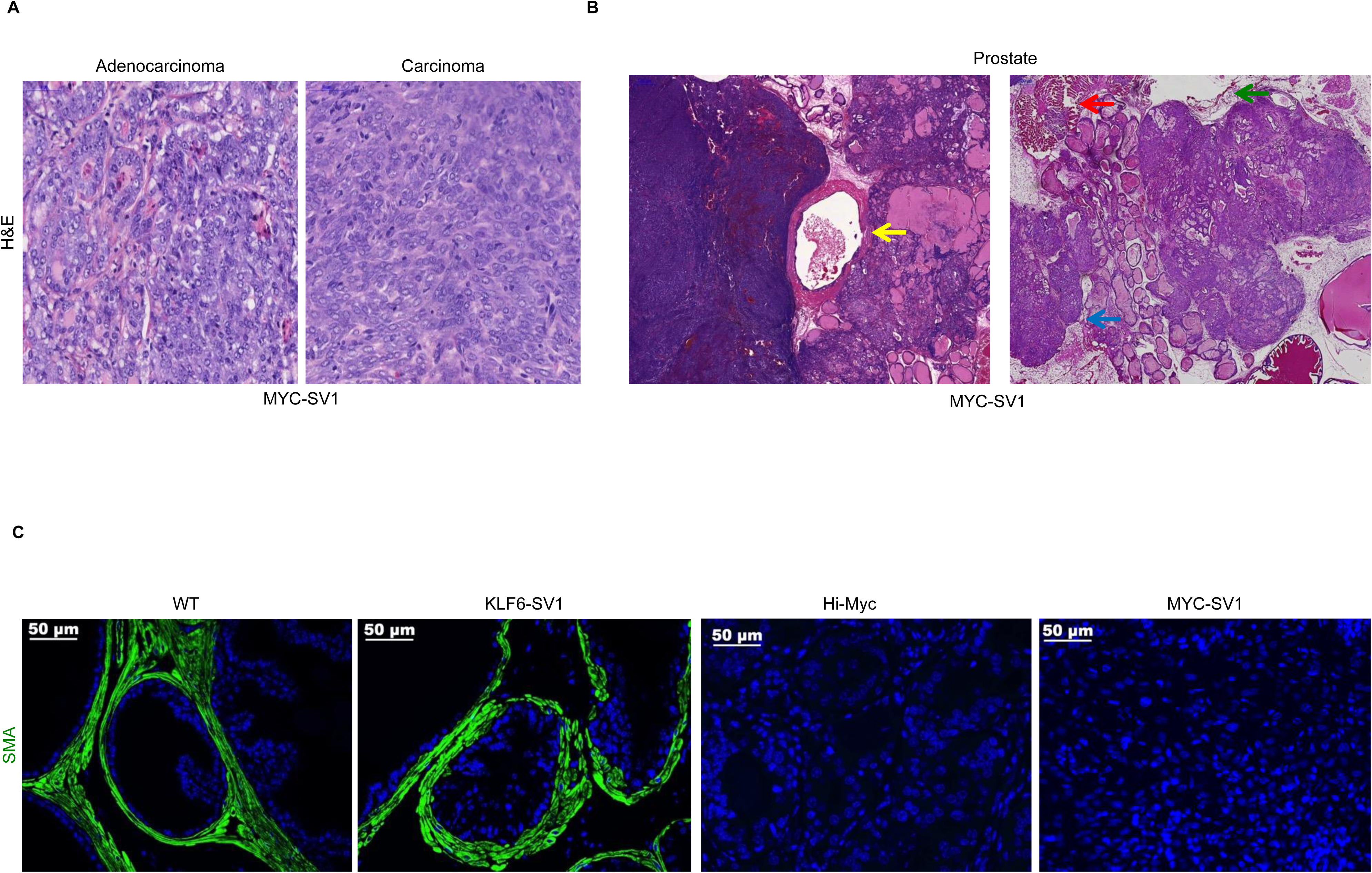
Related to Figure 1. MYC-SV1 tumors are poorly differentiated carcinomas and locally invasive. **A**, Hematoxylin and Eosin (H&E) staining of MYC-SV1 mouse prostate. **B,** An invasive adenocarcinoma lesion in a MYC-SV1 mouse exhibiting invasion into the bladder (yellow arrow), periprostatic adipose (blue arrow), muscle (orange arrow), and prostatic capsule (green arrow). **C,** Representative image of smooth muscle actin (SMA) staining (green fluorescence) and DAPI nuclear staining (blue fluorescence). Blood vessels serve as a positive internal control (10x, scale bar=100 µm).

**Figure S2.**
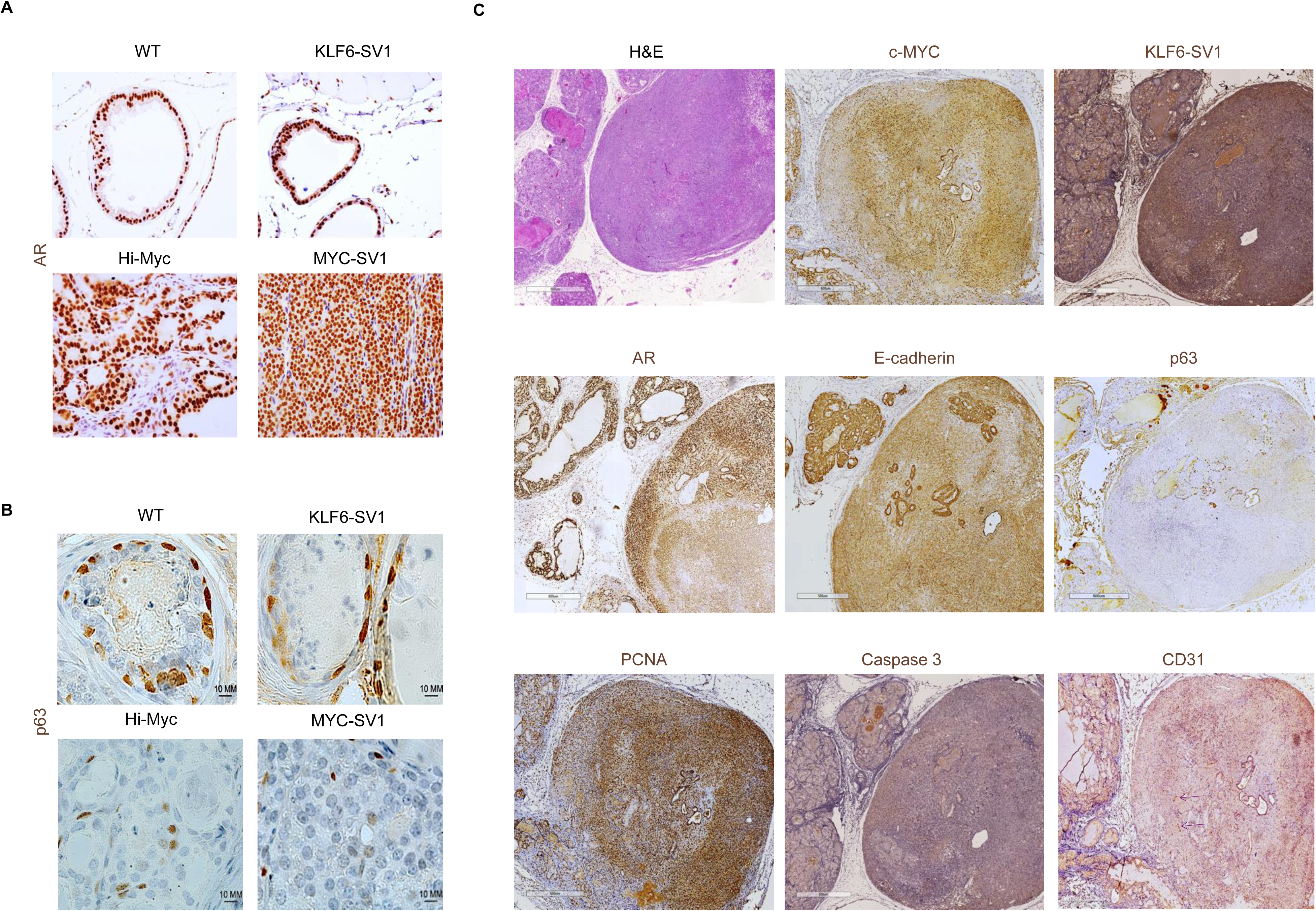
Related to Figure 2. Androgen and p63 expression in GEMMs. **A**, Strong nuclear AR is expressed in all mouse models. **B,** Loss of p63 is observed in tumors from MYC-SV1 mice. **C,** Molecular characterization by IHC staining with prostate lineage-marker specific antibodies on consecutive sections of a large poorly differentiated MYC-SV1 prostate carcinoma with un-involved well-differentiated adjacent adenocarcinoma (5x magnification, scale bar=600µm) for H&E, c-MYC, KLF6-SV1, AR, E-cadherin, p63, PCNA, Caspase-3, and CD31 with purple arrows point to blood vessels within the tumor.

**Figure S3.**
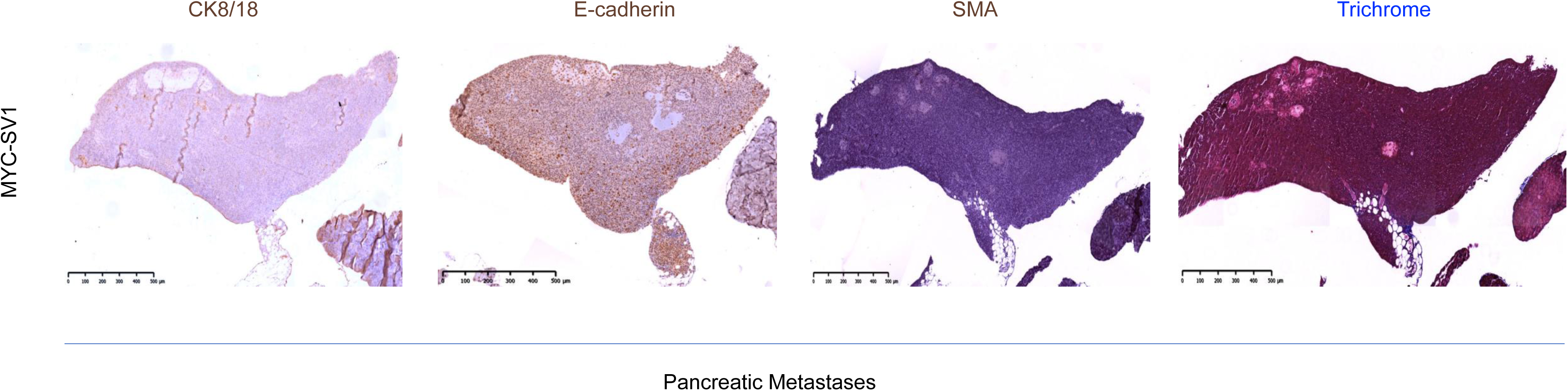
related to Figure 2. MYC-SV1 metastatic nodules are of epithelial origin. Expression of E-cadherin, Pan-CK, and alpha-smooth muscle actin.

**Figure S4.**
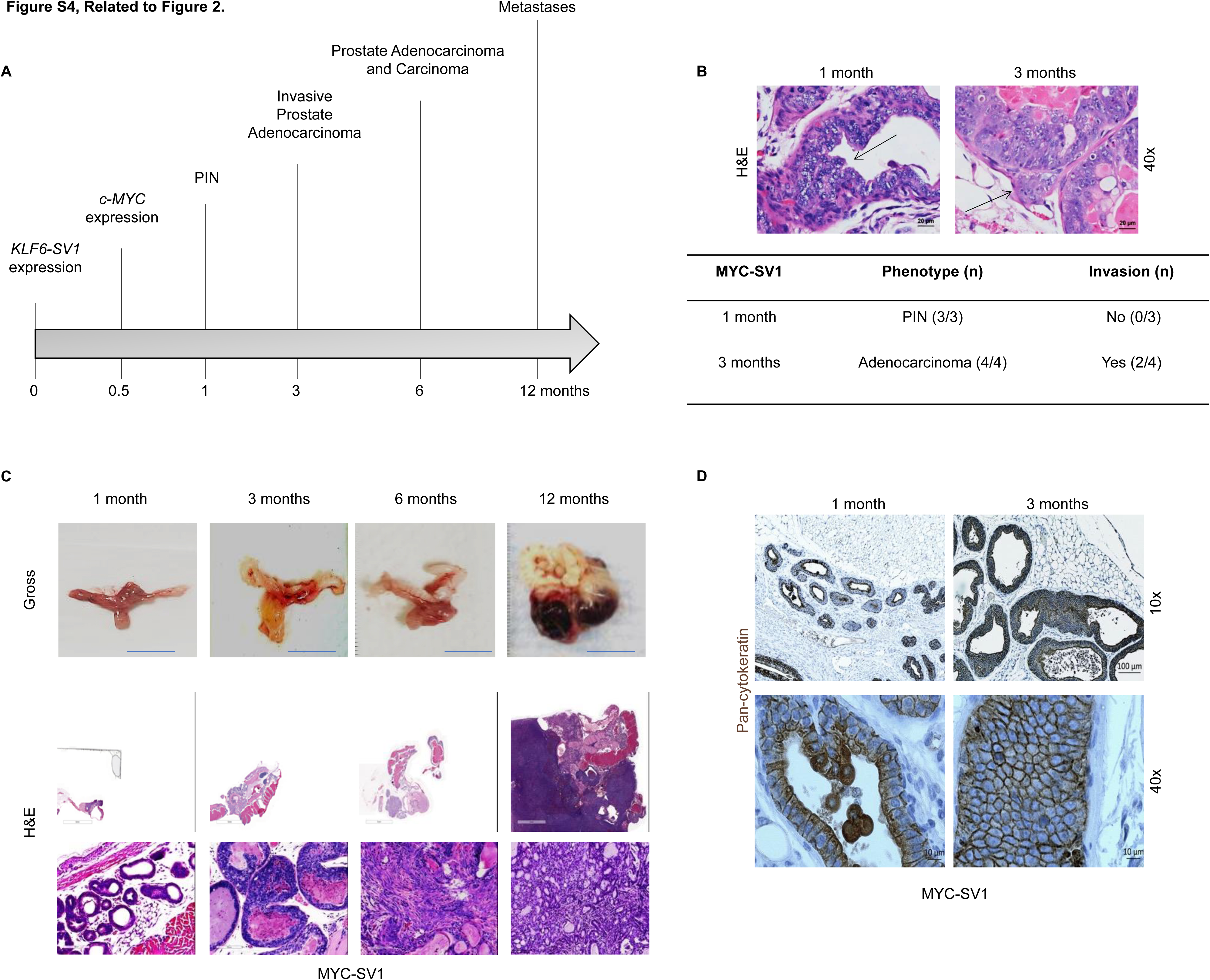
Related to Figure 1. MYC-SV1 tumor kinetics. **A**, Timeline of MYC-SV1 mPIN and prostate cancer development and progression. **B**, Gross and photomicrographs of FFPE sections stained with hematoxylin and eosin (H&E) of tumor development kinetics of the mouse urogenital system. Scale bar = 1cm. **C,** Early stages of tumor development in MYC-SV1 mice. mPIN (blue arrow); adenocarcinoma and invasion (black arrow). **D**, Expression of pan-cytokeratin.

**Figure S5.**
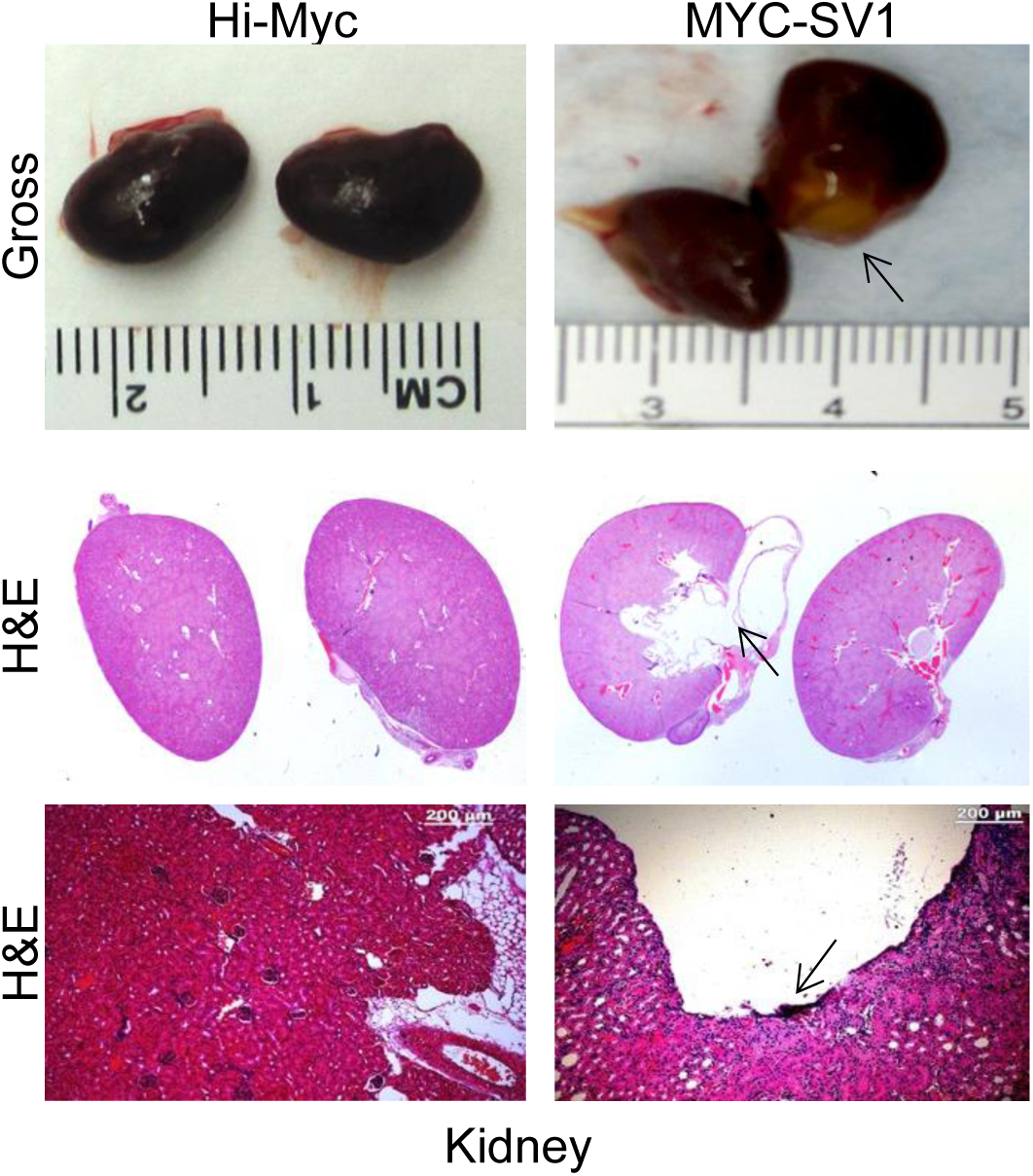
related to Figure 2. Histopathological presentation of hydronephrosis in MYC-SV1 mice. Representative gross and H&E-stained kidney sections from Hi-Myc and MYC-SV1 kidneys reveal hydronephrosis (arrow) in MYC-SV1 mice.

**Figure S6.**
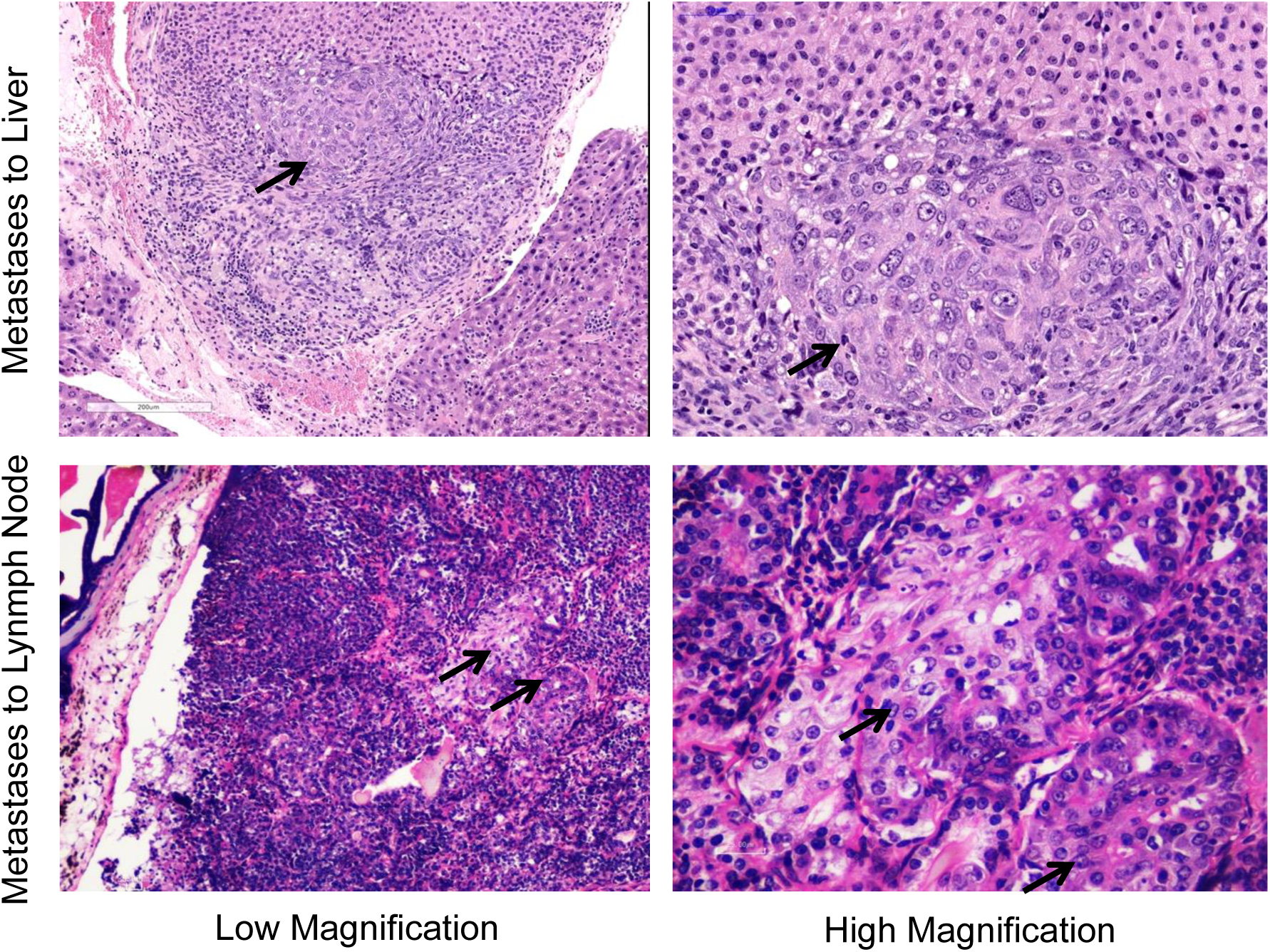
related to Figure 2. MYC-SV1 tumors metastasize to the liver and lymph node. Representative H&E stained FFPE sections of prostate cancer metastases to the liver and lymph nodes. Lack of smooth muscle actin and collagen expression (blue color) through Masson’s Trichrome staining confirmed that the metastatic foci were not of myoepithelial or fibroblast origin.

**Figure S7.**
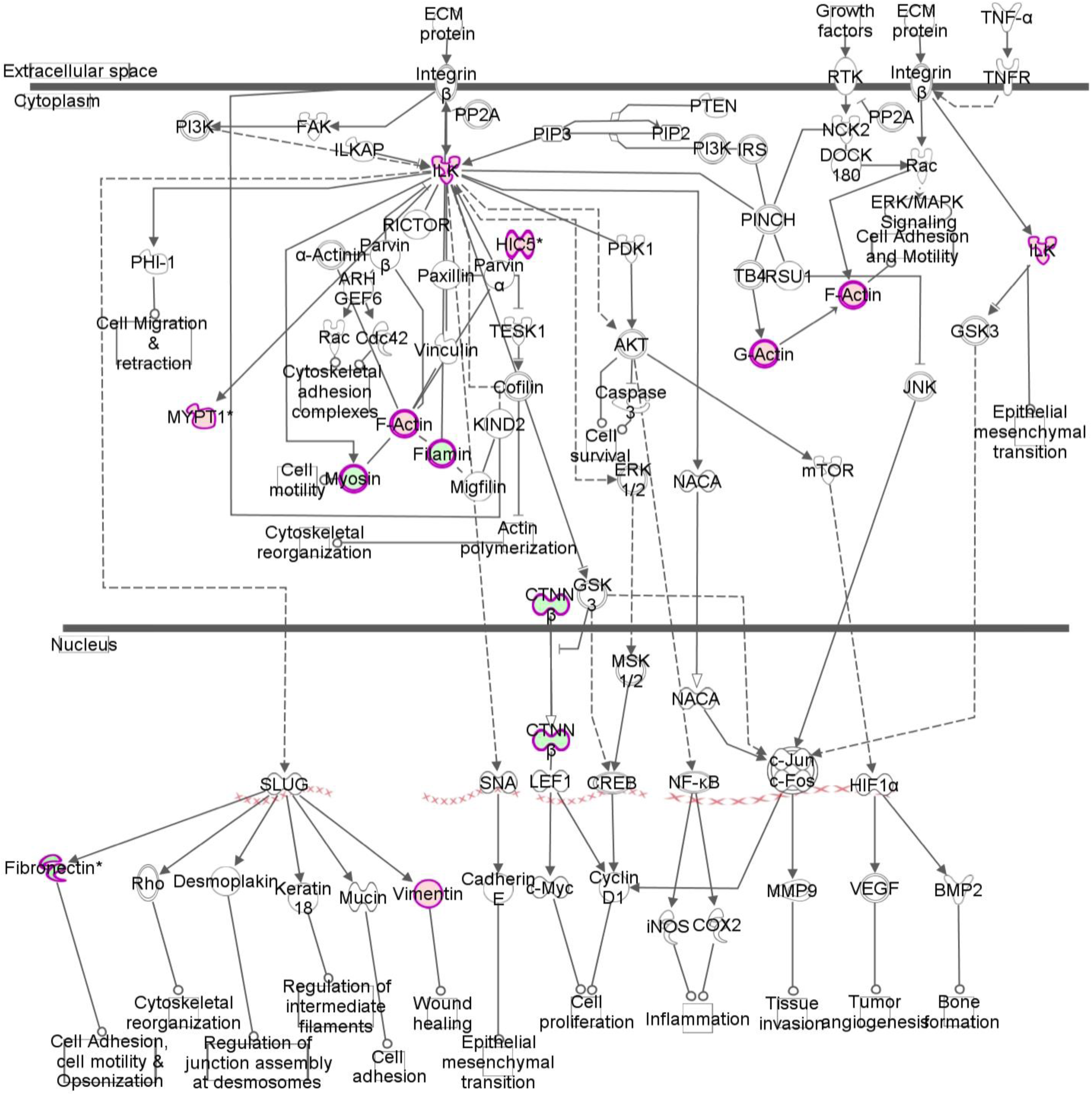
related to Figure 4. Ingenuity Pathway Analysis of Hi-Myc vs. MYC-SV1 tumors. Ingenuity Pathway Analysis (IPA) was utilized to elucidate the global implications of the differentially expressed proteins in our four mouse cohorts. The ILK pathway emerged as a top canonical pathway associated with our proteomics data. Changes in differential protein expression are depicted in red (upregulated) and green (downregulated). Proteins identified within this pathway included F-Actin, HIC5, and vimentin.

**Figure S8.**
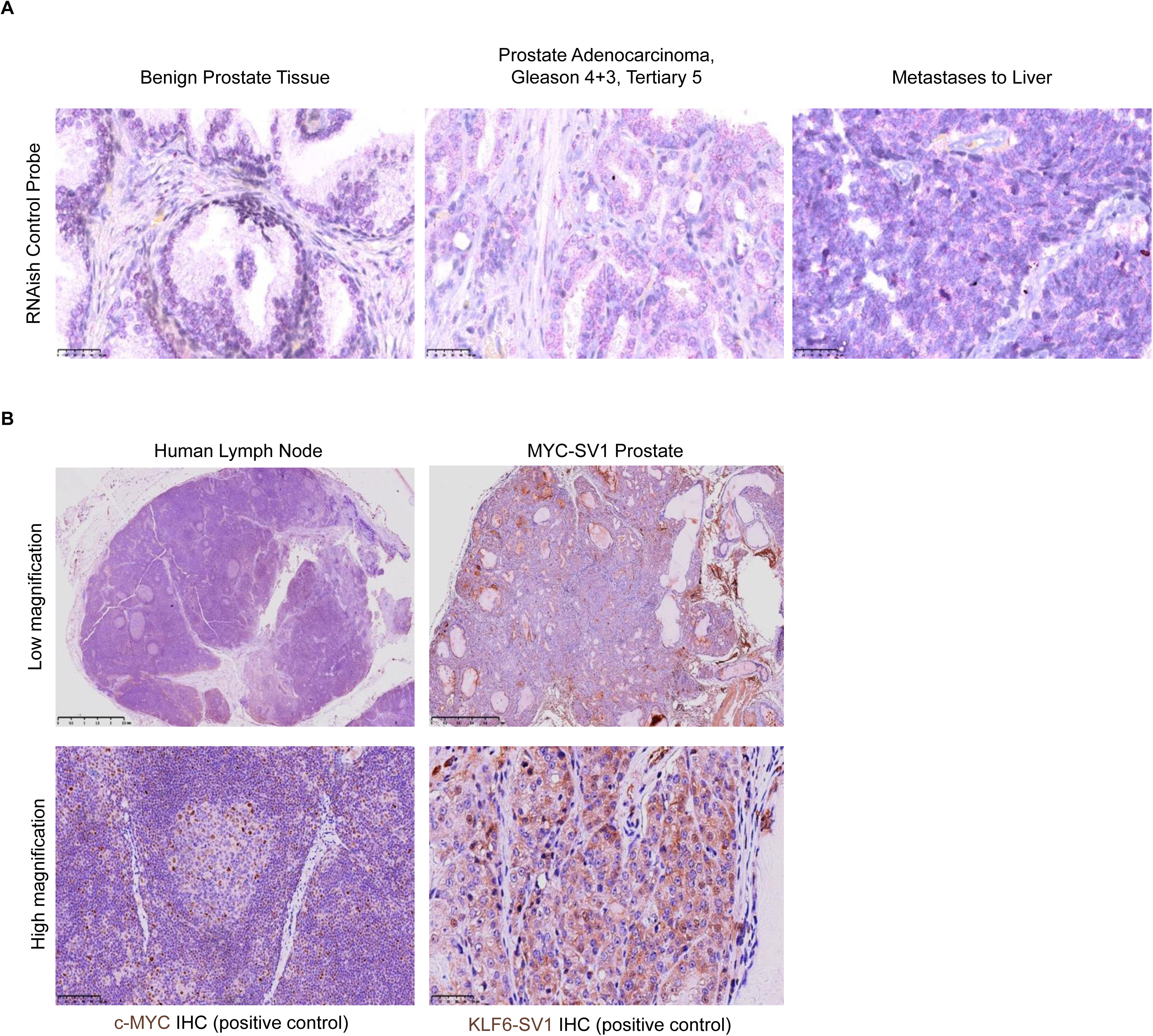
related to Figure 7. *RNAish* and IHC staining of positive control specimens for KLF6-SV1 and c-MYC. **A**, Human prostate tissue stained with control probes. **B,** Human lymph node tissue stained for c-MYC protein (left). MYC-SV1 mouse prostate stained for KLF6-SV1 protein (right).

## References

1. Siegel RL, Miller KD, Fuchs HE, Jemal A. Cancer statistics, 2021. CA Cancer J Clin. 2021 Jan;71(1):7–33.

2. Dhanasekaran SM, Barrette TR, Ghosh D, Shah R, Varambally S, Kurachi K, Pienta KJ, Rubin MA, Chinnaiyan AM. Delineation of prognostic biomarkers in prostate cancer. Nature. 2001 Aug 23;412(6849):822–6.

3. Lapointe J, Li C, Higgins JP, van de Rijn M, Bair E, Montgomery K, Ferrari M, Egevad L, Rayford W, Bergerheim U, Ekman P, DeMarzo AM, Tibshirani R, Botstein D, Brown PO, Brooks JD, Pollack JR. Gene expression profiling identifies clinically relevant subtypes of prostate cancer. Proc Natl Acad Sci U S A. 2004 Jan 20;101(3):811–6.

4. Taylor BS, Schultz N, Hieronymus H, Gopalan A, Xiao Y, Carver BS, Arora VK, Kaushik P, Cerami E, Reva B, Antipin Y, Mitsiades N, Landers T, Dolgalev I, Major JE, Wilson M, Socci ND, Lash AE, Heguy A, Eastham JA, Scher HI, Reuter VE, Scardino PT, Sander C, Sawyers CL, Gerald WL. Integrative genomic profiling of human prostate cancer. Cancer Cell. 2010 Jul 13;18(1):11–22.

5. Grasso CS, Wu YM, Robinson DR, Cao X, Dhanasekaran SM, Khan AP, Quist MJ, Jing X, Lonigro RJ, Brenner JC, Asangani IA, Ateeq B, Chun SY, Siddiqui J, Sam L, Anstett M, Mehra R, Prensner JR, Palanisamy N, Ryslik GA, Vandin F, Raphael BJ, Kunju LP, Rhodes DR, Pienta KJ, Chinnaiyan AM, Tomlins SA. The mutational landscape of lethal castration-resistant prostate cancer. Nature. 2012 Jul 12;487(7406):239–43.

6. Zhang X, Lee C, Ng PY, Rubin M, Shabsigh A, Buttyan R. Prostatic neoplasia in transgenic mice with prostate-directed overexpression of the c-myc oncoprotein. Prostate. 2000 Jun 1;43(4):278–85.

7. Ellwood-Yen K, Graeber TG, Wongvipat J, Iruela-Arispe ML, Zhang J, Matusik R, Thomas GV, Sawyers CL. Myc-driven murine prostate cancer shares molecular features with human prostate tumors. Cancer Cell. 2003 Sep;4(3):223–38.

8. Ellis L, Ku S, Li Q, Azabdaftari G, Seliski J, Olson B, Netherby CS, Tang DG, Abrams SI, Goodrich DW, Pili R. Generation of a C57BL/6 MYC-Driven Mouse Model and Cell Line of Prostate Cancer. Prostate. 2016 Sep;76(13):1192–202.

9. Kim J, Eltoum IE, Roh M, Wang J, Abdulkadir SA. Interactions between cells with distinct mutations in c-MYC and Pten in prostate cancer. PLoS Genet. 2009 Jul;5(7):e1000542.

10. Yang G, Goltsov AA, Ren C, Kurosaka S, Edamura K, Logothetis R, DeMayo FJ, Troncoso P, Blando J, DiGiovanni J, Thompson TC. Caveolin-1 upregulation contributes to c-Myc-induced high-grade prostatic intraepithelial neoplasia and prostate cancer. Mol Cancer Res. 2012 Feb;10(2):218–29.

11. Nguyen HG, Conn CS, Kye Y, Xue L, Forester CM, Cowan JE, Hsieh AC, Cunningham JT, Truillet C, Tameire F, Evans MJ, Evans CP, Yang JC, Hann B, Koumenis C, Walter P, Carroll PR, Ruggero D. Development of a stress response therapy targeting aggressive prostate cancer. Sci Transl Med. 2018 May 2;10(439).

12. Hubbard GK, Mutton LN, Khalili M, McMullin RP, Hicks JL, Bianchi-Frias D, Horn LA, Kulac I, Moubarek MS, Nelson PS, Yegnasubramanian S, De Marzo AM, Bieberich CJ. Combined MYC Activation and Pten Loss Are Sufficient to Create Genomic Instability and Lethal Metastatic Prostate Cancer. Cancer Res. 2016 Jan 15;76(2):283–92.

13. Clegg NJ, Couto SS, Wongvipat J, Hieronymus H, Carver BS, Taylor BS, Ellwood-Yen K, Gerald WL, Sander C, Sawyers CL. MYC cooperates with AKT in prostate tumorigenesis and alters sensitivity to mTOR inhibitors. PLoS One. 2011 Mar 4;6(3):e17449.

14. King JC, Xu J, Wongvipat J, Hieronymus H, Carver BS, Leung DH, Taylor BS, Sander C, Cardiff RD, Couto SS, Gerald WL, Sawyers CL. Cooperativity of TMPRSS2-ERG with PI3-kinase pathway activation in prostate oncogenesis. Nat Genet. 2009 May;41(5):524–6.

15. Jin RJ, Lho Y, Connelly L, Wang Y, Yu X, Saint Jean L, Case TC, Ellwood-Yen K, Sawyers CL, Bhowmick NA, Blackwell TS, Yull FE, Matusik RJ. The nuclear factor-kappaB pathway controls the progression of prostate cancer to androgen-independent growth. Cancer Res. 2008 Aug 15;68(16):6762–9.

16. Wang J, Kim J, Roh M, Franco OE, Hayward SW, Wills ML, Abdulkadir SA. Pim1 kinase synergizes with c-MYC to induce advanced prostate carcinoma. Oncogene. 2010 Apr 29;29(17):2477–87.

17. Nandana S, Ellwood-Yen K, Sawyers C, Wills M, Weidow B, Case T, Vasioukhin V, Matusik R. Hepsin cooperates with MYC in the progression of adenocarcinoma in a prostate cancer mouse model. Prostate. 2010 May 1;70(6):591–600.

18. Liu B, Gong S, Li Q, Chen X, Moore J, Suraneni MV, Badeaux MD, Jeter CR, Shen J, Mehmood R, Fan Q, Tang DG. Transgenic overexpression of NanogP8 in the mouse prostate is insufficient to initiate tumorigenesis but weakly promotes tumor development in the Hi-Myc mouse model. Oncotarget. 2017 Apr 18;8(32):52746–52760.

19. Narla G, Difeo A, Reeves HL, Schaid DJ, Hirshfeld J, Hod E, Katz A, Isaacs WB, Hebbring S, Komiya A, McDonnell SK, Wiley KE, Jacobsen SJ, Isaacs SD, Walsh PC, Zheng SL, Chang BL, Friedrichsen DM, Stanford JL, Ostrander EA, Chinnaiyan AM, Rubin MA, Xu J, Thibodeau SN, Friedman SL, Martignetti JA. A germline DNA polymorphism enhances alternative splicing of the KLF6 tumor suppressor gene and is associated with increased prostate cancer risk. Cancer Res. 2005 Feb 15;65(4):1213–22.

20. Narla G, DiFeo A, Fernandez Y, Dhanasekaran S, Huang F, Sangodkar J, Hod E, Leake D, Friedman SL, Hall SJ, Chinnaiyan AM, Gerald WL, Rubin MA, Martignetti JA. KLF6-SV1 overexpression accelerates human and mouse prostate cancer progression and metastasis. J Clin Invest. 2008 Aug;118(8):2711–21.

21. Narla G, DiFeo A, Yao S, Banno A, Hod E, Reeves HL, Qiao RF, Camacho-Vanegas O, Levine A, Kirschenbaum A, Chan AM, Friedman SL, Martignetti JA. Targeted inhibition of the KLF6 splice variant, KLF6 SV1, suppresses prostate cancer cell growth and spread. Cancer Res. 2005 Jul 1;65(13):5761–8.

22. Roh M, Kim J, Song C, Wills M, Abdulkadir SA. Transgenic mice for Cre-inducible overexpression of the oncogenes c-MYC and Pim-1 in multiple tissues. Genesis. 2006 Oct;44(10):447–53.

23. Watson PA, Ellwood-Yen K, King JC, Wongvipat J, Lebeau MM, Sawyers CL. Context-dependent hormone-refractory progression revealed through characterization of a novel murine prostate cancer cell line. Cancer Res. 2005 Dec 15;65(24):11565–71.

24. Podsypanina K, Politi K, Beverly LJ, Varmus HE. Oncogene cooperation in tumor maintenance and tumor recurrence in mouse mammary tumors induced by Myc and mutant Kras. Proc Natl Acad Sci U S A. 2008 Apr 1;105(13):5242–7.

25. Soucek L, Whitfield J, Martins CP, Finch AJ, Murphy DJ, Sodir NM, Karnezis AN, Swigart LB, Sergio Nasi S, Evan GI. Modelling Myc Inhibition as a Cancer Therapy. Nature. 2008 Oct 2;455(7213):679–83.

26. Shachaf CM, Kopelman AM, Arvanitis C, Karlsson A, Beer S, Mandl S, Bachmann MH, Borowsky AD, Ruebner B, Cardiff RD, Yang Q, Bishop JM, Contag CH, Felsher DW. MYC inactivation uncovers pluripotent differentiation and tumour dormancy in hepatocellular cancer. Nature. 2004 Oct 28;431(7012):1112–7.

27. Jain M, Arvanitis C, Chu K, Dewey W, Leonhardt E, Trinh M, Sundberg CD, Bishop JM, Felsher DW. Sustained loss of a neoplastic phenotype by brief inactivation of MYC. Science. 2002 Jul 5;297(5578):102–4.

28. Gurel B, Iwata T, Koh CM, Jenkins RB, Lan F, Van Dang C, Hicks JL, Morgan J, Cornish TC, Sutcliffe S, Isaacs WB, Luo J, De Marzo AM. Nuclear MYC protein overexpression is an early alteration in human prostate carcinogenesis. Mod Pathol. 2008 Sep;21(9):1156–67.

29. Beltran H, Prandi D, Mosquera JM, Benelli M, Puca L, Cyrta J, Marotz C, Giannopoulou E, Chakravarthi BV, Varambally S, Tomlins SA, Nanus DM, Tagawa ST, Van Allen EM, Elemento O, Sboner A, Garraway LA, Rubin MA, Demichelis F. Divergent clonal evolution of castration-resistant neuroendocrine prostate cancer. Nat. Med. 2016;22:298–305.

30. Das R, Gregory PA, Hollier BG, Tilley WD, Selth LA. Epithelial plasticity in prostate cancer: principles and clinical perspectives. Trends Mol Med. 2014 Nov;20(11):643–51.

31. Yang J, Antin P, Berx G, Blanpain C, Brabletz T, Bronner M, Campbell K, Cano A, Casanova J, Christofori G, Dedhar S, Derynck R, Ford HL, Fuxe J, García de Herreros A, Goodall GJ, Hadjantonakis AK, Huang RJY, Kalcheim C, Kalluri R, Kang Y, Khew-Goodall Y, Levine H, Liu J, Longmore GD, Mani SA, Massagué J, Mayor R, McClay D, Mostov KE, Newgreen DF, Nieto MA, Puisieux A, Runyan R, Savagner P, Stanger B, Stemmler MP, Takahashi Y, Takeichi M, Theveneau E, Thiery JP, Thompson EW, Weinberg RA, Williams ED, Xing J, Zhou BP, Sheng G; EMT International Association (TEMTIA). Guidelines and definitions for research on epithelial-mesenchymal transition. Nat Rev Mol Cell Biol. 2020 Apr 16.

32. Zhao Y, Yan Q, Long X, Chen X, Wang Y. Vimentin affects the mobility and invasiveness of prostate cancer cells. Cell Biochem Funct. 2008 Sep-Oct;26(5):571–7.

33. Armstrong AJ, Marengo MS, Oltean S, Kemeny G, Bitting RL, Turnbull JD, Herold CI, Marcom PK, George DJ, Garcia-Blanco MA. Circulating tumor cells from patients with advanced prostate and breast cancer display both epithelial and mesenchymal markers. Mol Cancer Res. 2011 Aug;9(8):997–1007.

34. Lang SH, Hyde C, Reid IN, Hitchcock IS, Hart CA, Gordon Bryden AA, Villette JM, Stower MJ, Maitland NJ. Enhanced expression of vimentin in motile prostate cell lines and in poorly differentiated and metastatic prostate carcinoma. Prostate. 2002 Sep 1;52(4):253–63.

35. Hanahan D, Weinberg RA. Hallmarks of cancer: the next generation. Cell. 2011 Mar 4;144(5):646–74.

36. Tomechko SE, Liu G, Tao M, Schlatzer D, Powell CT, Gupta S, Chance MR, Daneshgari F. Tissue specific dysregulated protein subnetworks in type 2 diabetic bladder urothelium and detrusor muscle. Mol Cell Proteomics. 2015 Mar;14(3):635–45.

37. Wisniewski JR, Zougman A, Nagaraj N, Mann M. Universal sample preparation method for proteome analysis. Nat Methods. 2009 May;6(5):359–62.

38. Mi H, Muruganujan A, Ebert D, Huang X, Thomas PD. PANTHER version 14: more genomes, a new PANTHER GO-slim and improvements in enrichment analysis tools. Nucleic Acids Res. 2019 Jan 8;47(D1):D419–D426.

39. Park K, Tomlins SA, Mudaliar KM, Chiu YL, Esgueva R, Mehra R, Suleman K, Varambally S, Brenner JC, MacDonald T, Srivastava A, Tewari AK, Sathyanarayana U, Nagy D, Pestano G, Kunju LP, Demichelis F, Chinnaiyan AM, Rubin MA. Antibody-based detection of ERG rearrangement-positive prostate cancer. Neoplasia. 2010 Jul;12(7):590–8.

40. Arriaga JM, Panja S, Alshalalfa M, Zhao J, Zou M, Giacobbe A, Madubata CJ, Yeji Kim J, Rodriguez A, Coleman I, Virk RK, Hibshoosh H, Ertunc O, Ozbek B, Fountain J, Karnes RJ, Luo J, Antonarakis ES, Nelson PS, Feng FY, Rubin MA, De Marzo AM, Rabadan R, Sims PA, Mitrofanova A, Abate-Shen C. A MYC and RAS co-activation signature in localized prostate cancer drives bone metastasis and castration resistance. Nat Cancer. 2020 Nov;1(11):1082–1096.

41. Gurel B, Iwata T, Koh CM, Jenkins RB, Lan F, Van Dang C, Hicks JL, Morgan J, Cornish TC, Sutcliffe S, Isaacs WB, Luo J, De Marzo AM. Nuclear MYC protein overexpression is an early alteration in human prostate carcinogenesis. Mod Pathol. 2008 Sep;21(9):1156–67.

42. Melis MHM, Nevedomskaya E, van Burgsteden J, Cioni B, van Zeeburg HJT, Song JY, Zevenhoven J, Hawinkels LJAC, de Visser KE, Bergman AM. The adaptive immune system promotes initiation of prostate carcinogenesis in a human c-Myc transgenic mouse model. Oncotarget. 2017 Sep 28;8(55):93867–93877.

43. Shappell SB, Thomas GV, Roberts RL, Herbert R, Ittmann MM, Rubin MA, Humphrey PA, Sundberg JP, Rozengurt N, Barrios R, Ward JM, Cardiff RD. Prostate pathology of genetically engineered mice: definitions and classification. The consensus report from the Bar Harbor meeting of the Mouse Models of Human Cancer Consortium Prostate Pathology Committee. Cancer Res. 2004 Mar 15;64(6):2270–305.

44. Thompson TC, Southgate J, Kitchener G, Land H. Multistage carcinogenesis induced by ras and myc oncogenes in a reconstituted organ. Cell. 1989 Mar 24;56(6):917–30.

